# ADAR1 Regulates Lipid Remodeling to Dictate Ferroptosis Sensitivity

**DOI:** 10.1101/2025.01.16.633410

**Authors:** Che-Pei Kung, Nick D. Terzich, Ma Xenia G. Ilagan, Michael J. Prinsen, Madhurima Kaushal, Raleigh D. Kladney, Jade H. Weber, Alex R. Mabry, Luisangely Soto Torres, Emily R. Bramel, Eric C. Freeman, Thwisha Sabloak, Sua Ryu, William M. Weber, Kyle A. Cottrell, Leonard B. Maggi, Leah P. Shriver, Gary J. Patti, Jason D. Weber

## Abstract

Triple-negative breast cancer (TNBC), defined by the lack of estrogen, progesterone, and HER2 receptor expressions, is aggressive and lacks targeted treatment options. Adenosine deaminase acting on RNA 1 (ADAR1) has been shown to contribute to TNBC tumorigenesis by modulating an innate immune response. However, little is known about its role in the regulation of metabolic fitness of TNBC. We tested the hypothesis that ferroptosis is an ADAR1-protected vulnerability by showing that ADAR1 knockdown sensitizes TNBC cells to ferroptosis inducers. Lipidomic analyses showed that ADAR1 loss increased the abundance of phospholipids enriched with polyunsaturated fatty acids (PUFA), of which peroxidation is the main driver of ferroptosis. Transcriptomic analyses discovered that the proto-oncogene MDM2 contributes to the lipid remodeling phenotype. A ferroptosis-focused drug screen identified FDA-approved cobimetinib as a drug-repurposing candidate to synergize with ADAR1 loss. This finding supports further basic, pre-clinical, and clinical studies to develop novel therapeutic strategies for TNBC through ADAR1-mediated metabolic regulation.

## Introduction

Despite the recent development of PARP inhibitors, immunotherapy, and antibody-drug conjugates, there remains a significant gap in targeted therapies for TNBC, contributing to its poor prognosis^1^. A multi-prong approach is underway to investigate the immunogenic, epigenetic, and metabolic landscapes of TNBC to identify points of vulnerability^2–4^.

One such vulnerability is ferroptosis, a distinctive form of programmed cell death that depends on iron-mediated peroxidation of long polyunsaturated fatty acids (PUFA) to break down cell membranes^5^. Compared to other BC subtypes, TNBC cells express lower levels of iron exporter ferroportin, resulting in an increase in labile iron pools^6^. TNBC cells have also been shown to increase the uptake of cystine, a precursor for the antioxidant glutathione, rendering them more vulnerable to inhibitions of the glutamate/cystine antiporter xCT and glutathione effector GPX4^7,8^. TNBC’s sensitivity to ferroptosis is also correlated with abundance of PUFA, regulated by enzymes contributing to PUFA biosynthesis, such as Acyl-CoA synthetase long-chain family member 1 and 4 (ACSL1/4), as well as fatty acid desaturases 1 and 2 (FADS1/2)^9–11^.

Adenosine deaminase acting on RNA 1 (ADAR1, encoded by *ADAR*) converts adenosine to inosine within cellular dsRNA, in a process known as A-to-I editing. Elevated expression of ADAR1 is prevalent in breast cancer, and ADAR1-mediated editing contributes to tumor progression and resistance to therapy in breast cancer, including TNBC^12,13^. We have previously reported a TNBC-centric ADAR1 dependency phenomenon, in which ADAR1 protects TNBC cells from translational stress and type I interferon activation predisposed by elevated levels of interferon stimulated gene signatures^14^. ADAR1-mediated regulation of the innate immune response has been observed in several types of cancer, and the loss of ADAR1 has been shown to sensitize certain cancers to immunotherapy^15–18^. Through its ability to prevent the cytotoxic inflammatory response, ADAR1 also protects cancer cells from other anti-tumorigenic stresses, such as necroptosis and epigenetic modulation^19,20^. However, few studies have investigated the ability of ADAR1 to regulate the metabolic fitness of cancer cells and their sensitivity to metabolic stress. Considering the role of ADAR1 in metabolic regulation and the importance of cancer metabolism, it is important to fill these knowledge gaps^21,22^.

In the current study, we demonstrate that ADAR1 protects TNBC cells from ferroptosis by regulating the expression of MDM2 and lipid profiles. Our observations indicate broader applications of ADAR1-targeting therapies against TNBC.

## Results

### ADAR1 protects TNBC cells from ferroptosis induction

To determine whether ADAR1 inhibits ferroptosis directly, ADAR1 was knocked down with a short hairpin RNA (shRNA) in the TNBC cell line MDA-MB-231 (**Figure S1A**). The level of malondialdehyde (MDA), a byproduct and marker of lipid peroxidation, was measured to show that cells with ADAR1 knockdown (ADAR1-deficient) did not display induced ferroptosis phenotype compared to ADAR1-intact cells treated with a non-targeting shRNA (shNT) (**Figure S1B**). Similarly, by staining for both the reduced and oxidized forms of the lipid peroxidation probe BODIPY^TM^ 581/591 C11, it was revealed that ADAR1 knockdown did not improve the cells’ ability to oxidize PUFA (**Figure S1C-D**). Cell viability assay was performed to show that reduced proliferation and viability of MDA-MB-231 cells resulting from ADAR1 knockdown could be partially rescued by an apoptosis inhibitor (Z-VAD-FMK) or an eIF2α inhibitor (ISRIB), but not by ferroptosis inhibitors Ferrostatin-1 or Liproxstatin-1 (**Figure S1E**). These results suggest that the reduction of ADAR1 does not overtly induce ferroptosis phenotypes.

ADAR1-intact and -deficient MDA-MB-231 cells were then tested to determine whether ADAR1 expression influences the sensitivity of cells to ferroptosis induction. Ferroptosis can be induced by the GPX4 inhibitor RSL3, whose effect can be rescued by the ferroptosis inhibitors Ferrostatin-1 or UAMC, but not Z-VAD-FMK or necroptosis inhibitor Necrostatin-1 (**Figure S1F**). ADAR1-deficient cells were found to be more sensitive to RSL3 at concentrations as low as 50nM (**Figure 1A**). Cell viability assay was performed to determine the IC_50_ (half-maximum inhibitory concentration) of RSL3, and ADAR1-deficient cells (IC_50_ = 0.11µM vs 2.9µM) were 27 times more sensitive compared to ADAR1-intact cells (**Figure 1B**). A similar level of sensitization was observed when using another GPX4 inhibitor, ML162 (IC_50_ = 0.08µM vs 2.27µM) (**Figure 1C**).

**Figure 1.**
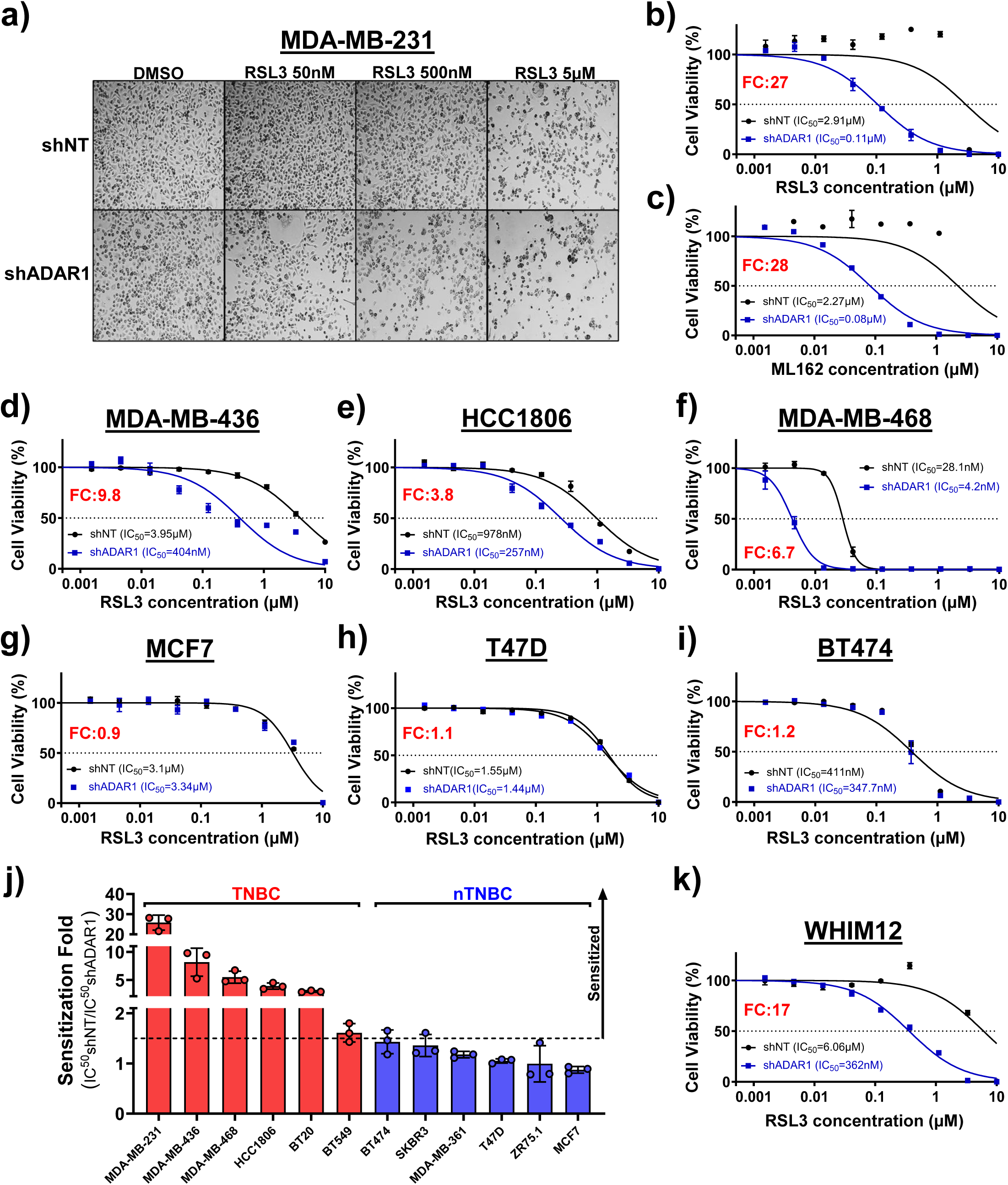
ADAR1 protects TNBC cells from ferroptosis induction **A)** Light micrographs of ADAR1-intact (shNT) and -deficient (shADAR1) MDA-MB-231 cells treated with RSL3 for 24h. Cell viability analyses were performed to determine IC_50_ values of ferroptosis inducers **B)** RSL3 and **C)** ML162 for MDA-MB-231 cells. IC_50_ of RSL3 were determined for **D)** MDA-MB-436, **E)** HCC1806, and **F)** MDA-MB-468 TNBC cells, as well as **G)** MCF7, **H)** T47D, and **I)** BT474 non-TNBC cells. Cell viability graphs are representatives of three replicates. **J)** Average levels of shADAR1-mediated sensitization to RSL3 among TNBC (red bars) and non-TNBC (nTNBC; blue bars) cell lines. Dashed line denotes an arbitrarily determined threshold (IC_50_shNT/IC_50_shADAR1 = 1.5) for notable shADAR1-mediated sensitization to RSL3. **K)** IC_50_ values of RSL3 for PDX-derived TNBC cell line WHIM12. FC (fold-change) represent levels of shADAR1-mediated sensitization to RSL3.

To evaluate whether this sensitization effect occurs in TNBC cells in addition to MDA-MB-231, multiple TNBC cell lines (MDA-MB-436, HCC1806, MDA-MB-468, BT20, BT549) were tested. In all TNBC cell lines tested, ADAR1 knockdown resulted in increased RSL3 sensitivity, ranging from 1.8-fold (BT549) to 9.8-fold (MDA-MB-436) (**Figure 1D-F, Figure S1G-H**). Multiple non-TNBC cell lines (MCF7, T47D, BT474, SKBR3, MDA-MB-361, ZR75.1) were also tested. Interestingly, by arbitrarily setting IC_50_shNT/IC_50_shADAR1=1.5 as the threshold for sensitization, none of the non-TNBC cell lines acquired higher sensitivity after ADAR1 knockdown (**Figure 1G-J, Figure S1I-K**). A cell line derived from a TNBC PDX (patient-derived xenograft), WHIM12, was also shown to be highly sensitized to RSL3 after ADAR1 knockdown (**Figure 1K**). These results indicate that ADAR1 protects TNBC, but not non-TNBC, cell lines from ferroptosis induction.

An orthotopic implantation mouse model was established to test whether these *in vitro* results can be recapitulated *in vivo*. MDA-MB-231 cells stably transduced with Tet-inducible shADAR1 were implanted in the mammary fat pad of female immunocompromised NSG mice (**Figure 2A-B**). Upon the formation of palpable tumors, half of the mice were switched to a diet containing doxycycline to induce ADAR1 knockdown (**Figure S2A**). Both control and doxycycline diet mice were subjected to intraperitoneal (i.p.) injection of RSL3 (100mg/kg weight; twice a week) for 5 weeks. Weekly health and weight monitoring did not suggest additional toxicity associated with RSL3 treatment (**Figure S2B**). Doxycycline diet or RSL3 treatment alone reduced tumor burdens, and mice that received both experienced significantly slower tumor progression (**Figure 1C-E, Figure S2C-E**). Immunohistochemical (IHC) staining of ADAR1 in resected tumors confirmed a sustained overall reduction in ADAR1 expression (**Figure 2F and S2F**). IHC staining of 4-Hydroxynonenal (4HNE), a product of lipid peroxidation, showed that the combination treatment enhanced the ferroptosis phenotype (**Figure 2F-G**). Moreover, showing a mosaic pattern of staining in doxycycline/RSL3-treated tumors, ADAR1 levels can be seen to be inversely correlated with 4HNE staining (**Figure S2G**). Together, these results suggest that ADAR1 reduction also sensitizes cancer cells to ferroptosis induction to suppress tumor progression *in vivo*.

**Figure 2:**
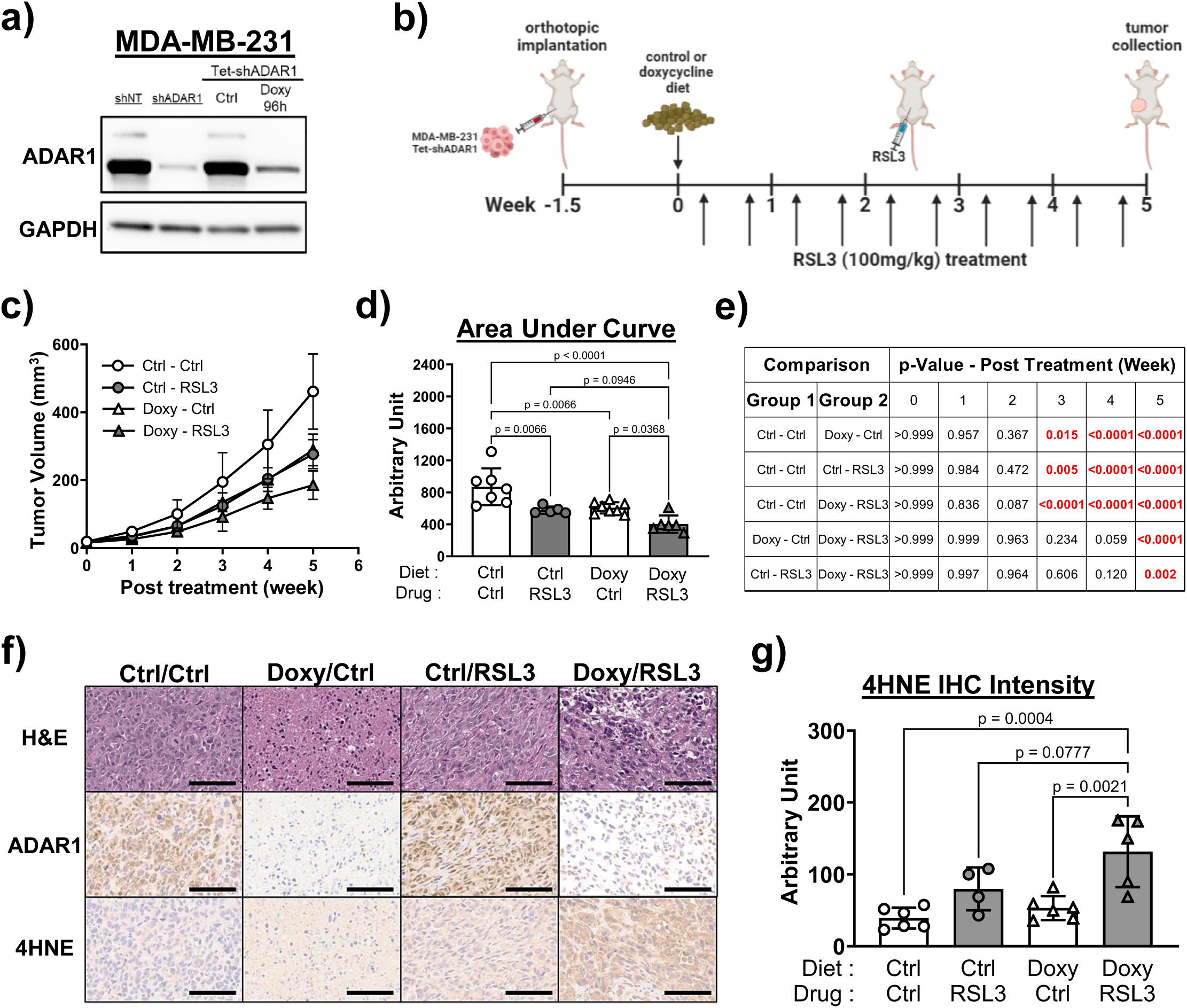
Combining ADAR1 reduction with RSL3 suppresses orthotopically implanted MDA-MB-231 xenograft progression *in vivo* **A)** Immunoblot analysis showed similar levels of ADAR1 knockdown in doxycycline-treated MDA-MB-231 cells with Tet-inducible shADAR1 compared to cells treated with shADAR1-containing lentivirus. Images are representative of three replicates. GAPDH, loading control. **B)** MDA-MB-231 cells (1.5 × 10^5^ per mouse) with Tet-inducible shADAR1 were implanted into mammary fat pads of female NSG mice. Ten days post implantation, half of the mice were switched to doxycycline-containing (625mg/kg) diet, followed by RSL3 treatments (100mg/kg weight; i.p. injection twice a week) for 5 weeks. **C)** Progression of overall tumor volumes, n=5-9 each group. Error bars mark standard deviation. **D)** Quantification of areas under curve (AUC) from C). Error bars mark standard deviation. **E)** Two-way ANOVA analysis to compare weekly measured overall tumor volumes. Statistically significant p-values (<0.05) were highlighted in red. **F)** Representative images of IHC staining of H&E, ADAR1 (p150 isoform), and 4HNE in resected tumor samples. Scale bar, 200µm. **G)** Quantification of IHC staining intensity of 4HNE. Error bars mark standard deviation. N=4-6 each group.

### Knockdown of ADAR1 leads to lipid remodeling

Multiple factors contribute to ferroptosis sensitivity, including ferrous iron availability, glutathione activity, and the relative abundance of PUFA over monounsaturated fatty acid (MUFA)^5^ (**Figure 3A**). To test the hypothesis that ADAR1-deficient cells are primed for ferroptosis induction, these elements were evaluated using the MDA-MB-231 cell line, as it is highly sensitive to ferroptosis induction by ADAR1 knockdown.

**Figure 3:**
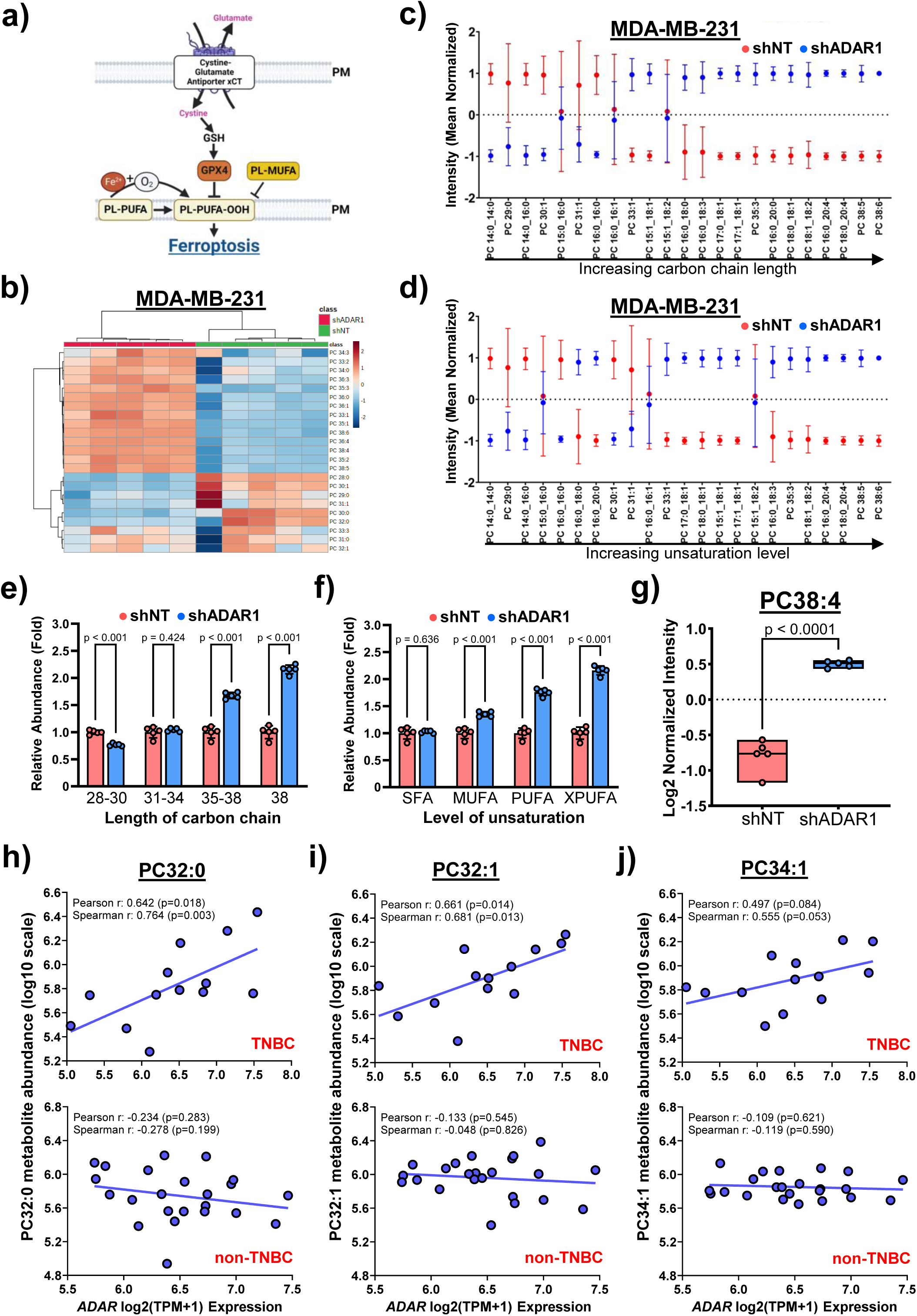
ADAR1 reduction leads to lipid remodeling **A)** Key regulators of ferroptosis. GSH, Glutathione. PL, phospholipid. MUFA, monounsaturated fatty acid. PUFA, polyunsaturated fatty acid. PM, plasma membrane. **B)** Heatmap of lipidomic analysis in MDA-MB-231 cells upon ADAR1 knockdown. PC, phosphatidylcholines. **C)** Relative abundance of PC species listed in the order of carbon chain length or **D)** lipid unsaturation. Values were log10 transformed, mean centered, and shown as mean over standard deviation ratio. n=5. Red dots denote ADAR1-intact, and blue dots denote ADAR1-deficient samples. **E)** Enrichment of PC species with different carbon chain lengths and **F)** different unsaturation levels (SFA, 0 double-bond; MUFA, 1 double-bond; PUFA, ≥ 2 double-bond; XPUFA, ≥ 4 double-bond) upon ADAR1 knockdown. Data from individual samples (n=4) were normalized to the average intensity in shNT samples. **G)** Relative abundance of PC38:4 between ADAR1-intact and -deficient MDA-MB-231 cells. Data from individual samples (n=5) were divided by the overall average of PC38:4 intensity in all samples and log2 transformed. Abundance of **H)** PC32:0, **I)** PC32:1, and **J)** PC34:1 positively correlate with *ADAR* (encoding ADAR1) expression level in TNBC (n=13), but not non-TNBC (n=23), cell lines. Data was extracted from metabolomics data set from DepMap (depmap.com).

In ADAR1-deficient cells, both the level of total glutathione and the ratio between reduced and total glutathione were unchanged (**Figure S3A-B**). FerroOrange staining was used to show that ADAR1-deficient cells do not display significantly higher levels of ferrous iron (**Figure S3C-D**). This suggests that elevated sensitivity to ferroptosis after ADAR1 knockdown cannot be attributed to altered glutathione activity or the abundance of ferrous iron.

Consequently, a targeted lipidomic analysis was performed to identify possible changes in lipid profiles after ADAR1 knockdown. Interestingly, a clear pattern change was detected with the phosphatidylcholine (PC) species, the most abundant phospholipid in mammalian cells and the most impactful phospholipid in controlling sensitivity to ferroptosis (**Figure 3B**). The observed lipid reprogramming disproportionally enriched PC species with longer carbon chains and higher degrees of unsaturation (**Figure 3C-F**). Some of the longest and most highly unsaturated PC species, such as PC38:4, are among the most significantly enriched PCs in ADAR1-deficient cells (**Figure 3G, Figure S3E**).

To investigate whether similar lipid reprogramming occurs in TNBC cells sensitized to ADAR1 knockdown to a lesser extent, lipidomic analysis was performed with ADAR1-deficient MDA-MB-436 cells (**Figure S3F**). Several highly unsaturated PCs, such as PC 34:3, 35:3, and 44:6, were enriched after ADAR1 knockdown (**Figure S3G-I**). However, enrichment was not limited to PUFA, as some MUFA PC species (i.e. PC36:1, 37:1, 38:1) were also upregulated. Some PUFA PC species appeared to be reduced (**Figure S3G**).

Interestingly, PUFAs from other phospholipid families, such as phosphatidylserine (PS) and phosphatidylethanolamine (PE), were also enriched in ADAR1-deficient MDA-MB-436 cells (**Figure S3J-N**). These results showed that the reduction of ADAR1 leads to lipid remodeling, specifically enrichment of PUFA over MUFA and saturated fatty acid (SFA). The publicly available metabolomics dataset shows that ADAR1 expression levels were strongly correlated with the abundance of SFA (PC32:0) and MUFA (PC32:1 and PC34:1) species in TNBC, but not non-TNBC cells (**Figure 3H-J**). These data support the hypothesis that in TNBC cells, ADAR1 helps maintain FA homeostasis to protect cells from ferroptosis.

### ADAR1 regulates MDM2 expression to dictate lipid profiles and ferroptosis sensitivity

To identify mechanisms contributing to ADAR1-regulated lipid remodeling, four TNBC cell lines, including those that are more highly (>5-fold - MDA-MB-231, MDA-MB-468) or moderately (<5-fold - HCC1806, BT549) sensitized to RSL3 after ADAR1 knockdown, were subjected to RNA-seq analyses to compare transcriptomic profiles between ADAR1-intact and ADAR1-deficient cells (**Figure 4A, Table S1**).

**Figure 4:**
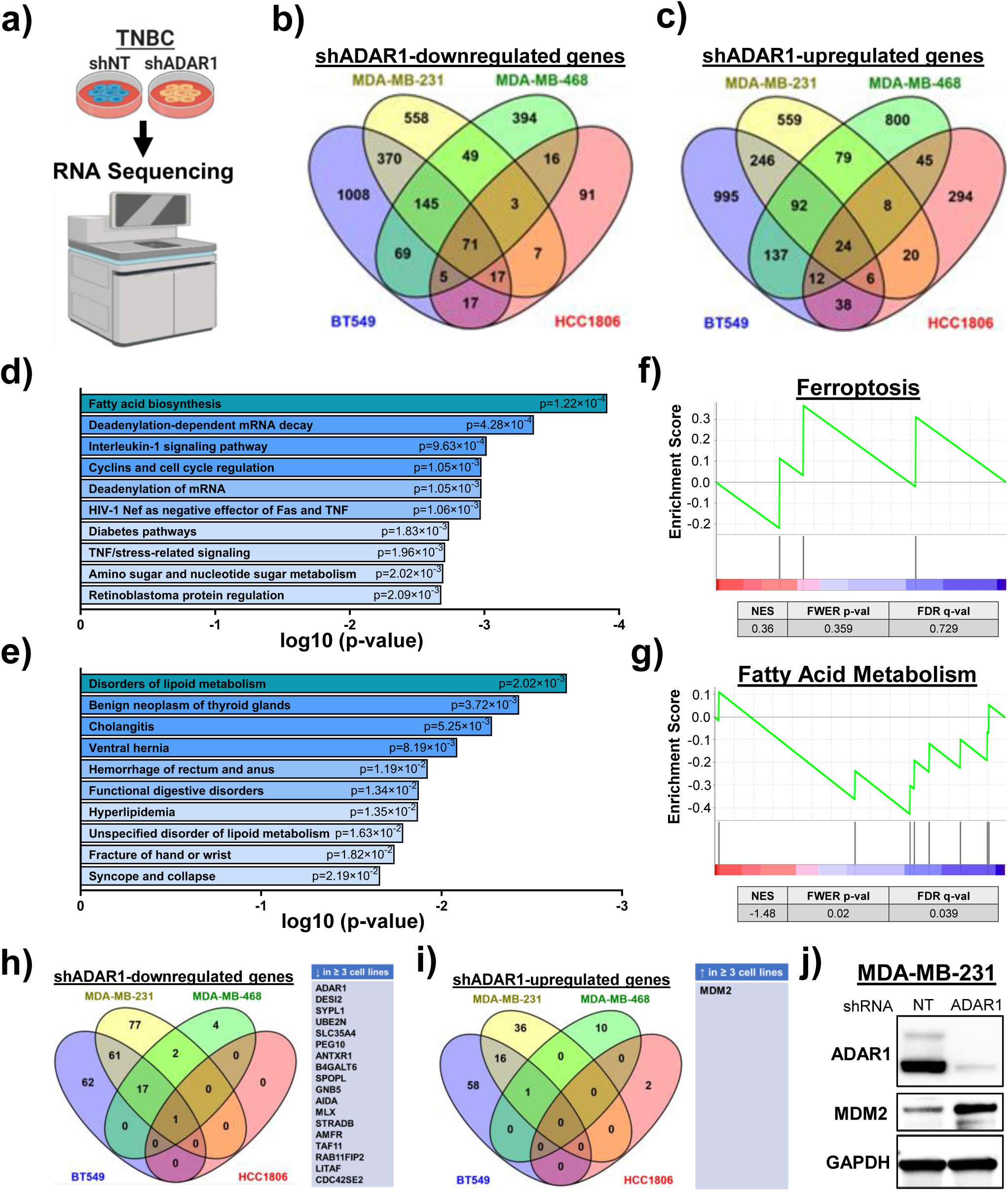
Transcriptomic regulation in TNBC cells upon ADAR1 knockdown **A)** RNA sequencing to identify significantly altered genes in ADAR1-deficient TNBC cells. **B)** Downregulated and **C)** upregulated genes in ADAR1-deficient TNBC cell lines. **D)** Significantly altered signaling pathways (BioPlanet_2019) and **E)** disease-related biological functions (PheWeb_2019) in ADAR1-deficient TNBC cells. Genes altered in at least 3 cell lines are included. **F)** GSEA enrichment plot of ferroptosis and **G)** fatty acid metabolism genes in ADAR1-deficient MDA-MB-231 cells. **H)** The most significantly downregulated and **I)** upregulated genes (FDR<0.01, Log2FC<-0.5 or >0.5) in ADAR1-deficient TNBC cells. Genes altered in at least 3 cell lines are listed on the right. **I)** Immunoblot analysis showed MDM2 induction upon ADAR1 knockdown in MDA-MB-231 cells. Images are representative of three replicates. GAPDH, loading control.

Pathway analyses were performed to provide a glimpse of cellular functions impacted by ADAR1 knockdown. The genes altered in at least three cell lines were from a variety of pathways, including metabolic regulations in cancer cells and lipid and fatty acid metabolism (**Figure 4B-E, Figure S4A-B, Table S2-4**). Gene set enrichment analysis (GSEA) with RNA-sequencing data from MDA-MB-231 cells showed that the reduction in ADAR1 resulted in significant changes in transcriptomes involved in fatty acid metabolism, but not in the ROS (reactive oxygen species) pathway, iron uptake, or direct ferroptosis regulation (**Figure 4F-G, Figure S4C-D**). GSEA results also suggest a lack of clear effect on interferon signaling or cell cycle regulation (**Figure S4E-F**). Interestingly, when other metabolic pathways were evaluated, ADAR1 knockdown had no effect on the glycolysis gene set but significantly altered the transcriptome involved in amino acid metabolism, suggesting that ADAR1 might affect the metabolic state of cancer cells beyond lipid programming (**Figure S4G-H**).

To narrow down the targets for further investigation, more stringent cutoff points (FDR<0.01, Log2FC>0.5 or <-0.5) were applied to identify the individual genes that were most easily altered. Seventeen genes (besides *ADAR*) were downregulated in at least 3 TNBC cell lines after ADAR1 knockdown, while only one (*MDM2*) was upregulated (**Figure 4H-I**). The induction of MDM2 protein in ADAR1-deficient cells was confirmed in multiple TNBC cell line models used in this study, including PDX-derived WHIM12 cells (**Figure 4J and S4I).**

The transcription of *MDM2* is known to be regulated by ADAR1-mediated editing in 3’UTR^23,24^. In ADAR1-deficient MDA-MB-231 and MDA-MB-468 cells, reduced levels of A-to-I/G editing were observed at the first reported editing site using both Sanger sequencing and RNA editing site-specific quantitative PCR (RESS-qPCR) (**Figure 5A-D**). Multiple studies have suggested that ADAR1-mediated RNA editing in *MDM2* 3’UTR can occur within a cluster of nearby editing sites, potentially representing a delicate machinery for MDM2 regulation^23,25,26^. By analyzing Sanger sequencing results from ADAR1-intact and -deficient MDA-MB-231 and MDA-MB-468 cells, it was revealed that despite both showing induced MDM2 expression after ADAR1 knockdown, the two cell lines displayed similar yet nonidentical editing profiles that are sensitive to ADAR1 knockdown (**Figure S5A-B**).

**Figure 5:**
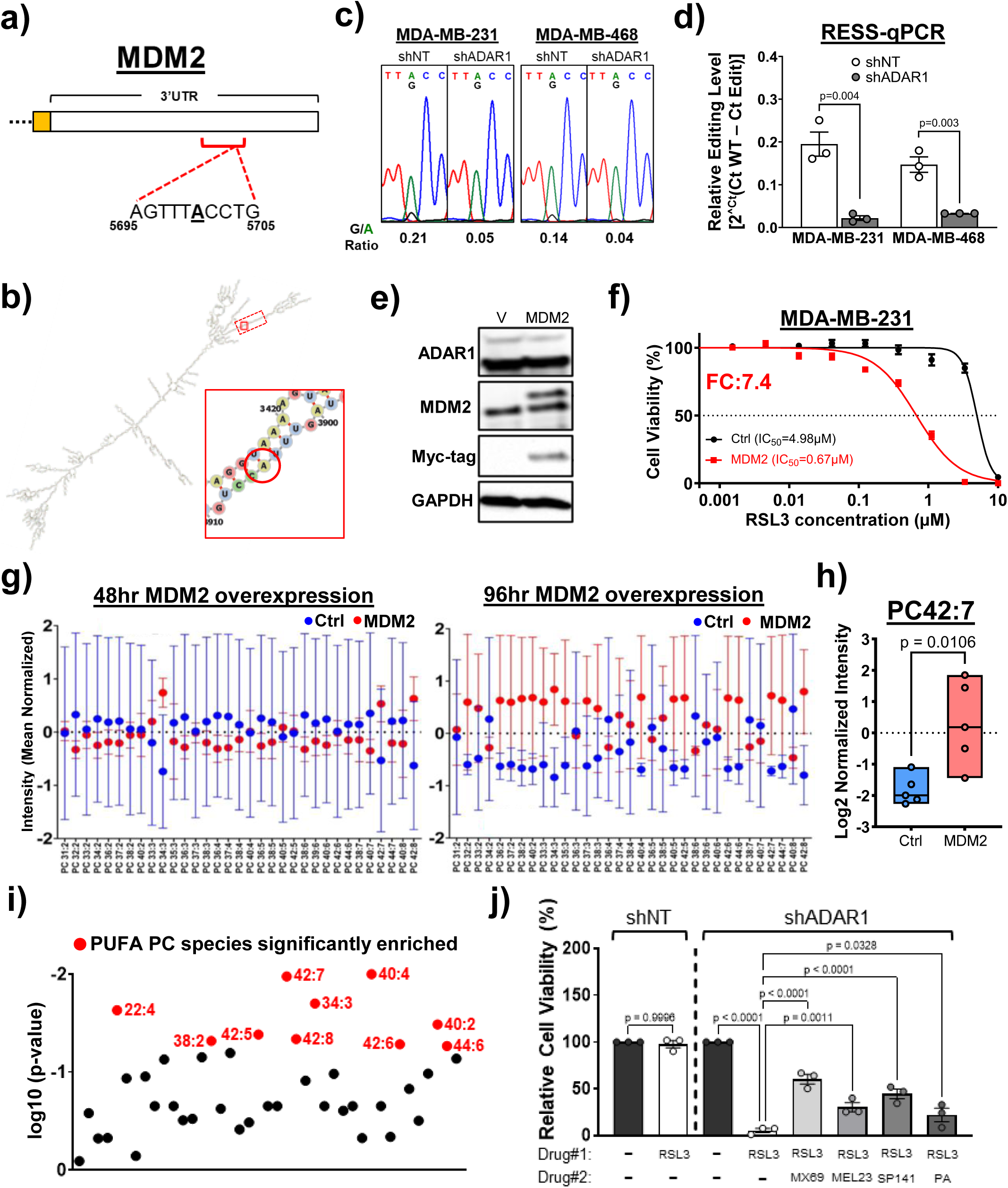
ADAR1 regulates MDM2 expression to dictate ferroptosis sensitivity **A)** Reported ADAR1 editing motifs (red bracket) and a representative site (bolded and underlined) in MDM2 3’UTR. Numbers mark the location within the open reading frame. **B)** Predicted RNA secondary structure of the entire MDM2 3’UTR and the reported ADAR1 editing motifs (dashed red box) and a specific site (circled in red in the zoomed-in square). **C)** Reduced editing of the representative MDM2 site in ADAR1-deficient TNBC cells. A (green) and G (black) reading represent non-edited and edited RNA, respectively. Ratio of G/A peak heights indicates editing levels. Sequence chromatograms shown are representative of 3 replicates. **D)** RESS-qPCR showed reduced editing of the MDM2 3’UTR site in ADAR1-deficient cells. n=3. Error bars mark standard error. **E)** Immunoblot analysis showed MDM2 (Myc-tagged) overexpression in MDA-MB-231 cells. GAPDH, loading control. **F)** IC_50_ of RSL3 for MDA-MB-231 cells with MDM2 overexpression. Graph is representative of three replicates. FC, fold-change of MDM2-mediated sensitization (IC_50_Control/IC_50_MDM2) to RSL3. **G)** Relative abundance of PUFA PC species, listed in the order of lipid unsaturation, in MDA-MB-231 cells with MDM2 overexpression (red dots) for 48h (left panel) and 96 h (right panel). n=5. **H)** Relative abundance of PC42:7 between control (Ctrl) and MDM2-overexpressing MDA-MB-231 cells. n=5. **I)** Significantly enriched PUFA PC species (red dots) in MDM2-overexpressing (96h) MDA-MB-231 cells. **J)** Relative cell viability within ADAR1-intact and -deficient MDA-MB-231 cells treated with RSL3 (100nM), as well as ADAR1-deficient cells co-treated with MDM2 inhibitors MX69 (10µM), MEL23 (10µM), and SP-141 (10µM), or PPAR-α agonist pirinixic acid (PA; 10µM) for 72h. n=3. Error bars mark standard error.

MDM2 has been shown to promote ferroptosis sensitivity by mediating lipid remodeling, including induction of PUFA levels^27^. To determine whether MDM2 is also capable of promoting ferroptosis sensitivity in TNBC cells, MDA-MB-231 cells overexpressing MDM2 were subjected to treatment with RSL3 (**Figure 5E**). Overexpression of MDM2 sensitized MDA-MB-231 cells to RSL3, albeit to a lesser extent compared to ADAR1 knockdown (**Figure 5F**). In particular, MDM2 overexpression resulted in induced sensitivity to RSL3 without affecting the proliferation potential of MDA-MB-231 cells, suggesting that MDM2 plays a ferroptosis-predisposing role in TNBC cells (**Figure S5C**).

To determine whether MDM2 promotes PUFA enrichment in TNBC cells similarly to ADAR1 knockdown, MDA-MB-231 cells possessing Tet-inducible MDM2 were subjected to lipidomic analysis to identify PC PUFA profiles 48 and 96 hours after doxycycline treatment (**Figure S5D**). At 48 hours, some PC PUFAs (i.e. PC34:3) were enriched. After 96 hours, a wider enrichment profile emerged, including inductions of several highly unsaturated PC species, such as PC42:7 and PC42:8 (**Figure 5G-I**). To confirm whether elevated ferroptosis sensitivity after ADAR1 reduction can be attributed to MDM2 induction, a rescue experiment was conducted with MDA-MB-231 cells treated with RSL3. The inflection point concentration of RSL3 (100nM) that distinguishes between ADAR1-intact and -deficient cells was used in combination with MDM2 inhibitors. All three MDM2 inhibitors used (MX69, MEL23, SP141) partially rescued ADAR1-deficient cells treated with RSL3 (**Figure 5J**). MDM2 was previously shown to promote ferroptosis sensitivity by suppressing the activity of peroxisome proliferator-activated receptor alpha (PPAR-α)^27^. Consistent with the report, an agonist of PPAR-α, pirinixic acid, was shown to confer a mild but significant rescue effect. These results indicate that ADAR1 regulates lipid profiles through MDM2 to protect TNBC cells from ferroptosis induction.

These results pointed to a paradoxical, tumor-suppressive role in TNBC for MDM2, commonly known as a proto-oncogene^28^. In fact, BC patients with the basal subtype (ESR1-/HER2-) with higher levels of tumor MDM2 expression have better survival outcomes (**Figure S5E**). This MDM2-associated survival advantage is amplified (HR = 0.61 VS 1.25) in patients that received systematic treatments (i.e. chemotherapy), potentially reflecting the fact that many existing therapies, such as platinum-based agents, induce ferroptosis^29,30^ (**Figure S5F-G**). One hypothesis is that instead of exerting its pro-tumorigenic activity through inhibiting wild-type p53, MDM2 engages in more p53-independnet functions such as lipid-remodeling/ferroptosis in TNBC cells that disproportionally possess deactivating p53 mutations that abrogate MDM2 interactions^31^. Supporting this hypothesis, high expression of MDM2 correlate strongly with better survival (HR = 0.39) in patients of the basal subtype with confirmed p53 mutations (**Figure S5H**).

### ADAR1 reduction synergizes with cobimetinib to suppress tumorigenesis

The most widely used ferroptosis inducers in preclinical studies, such as RSL3, remain distant from clinical applications due to poor pharmacological profiles and excessive toxicity^32^. Meanwhile, many drugs currently used in clinics or tested in clinical trials have ferroptosis-inducing activities^29^. Therefore, there is an opportunity to harness the lipid-remodeling effect of ADAR1 inhibition to enhance the efficacy of treatments readily available to patients.

To identify drug candidates that synergize with ADAR1 reduction, a high-throughput screen was performed to assess the effect of ADAR1 status on the sensitivity of MDA-MB-231 cells to drugs included in a ferroptosis-focused library (**Figure 6A**). Drugs that synergize with ADAR1 reduction (shADAR1-sensitized) were expected to suppress ADAR1-deficient cells more effectively, similarly to RSL3, which was used as a positive control (**Figure 6B-C**).

**Figure 6:**
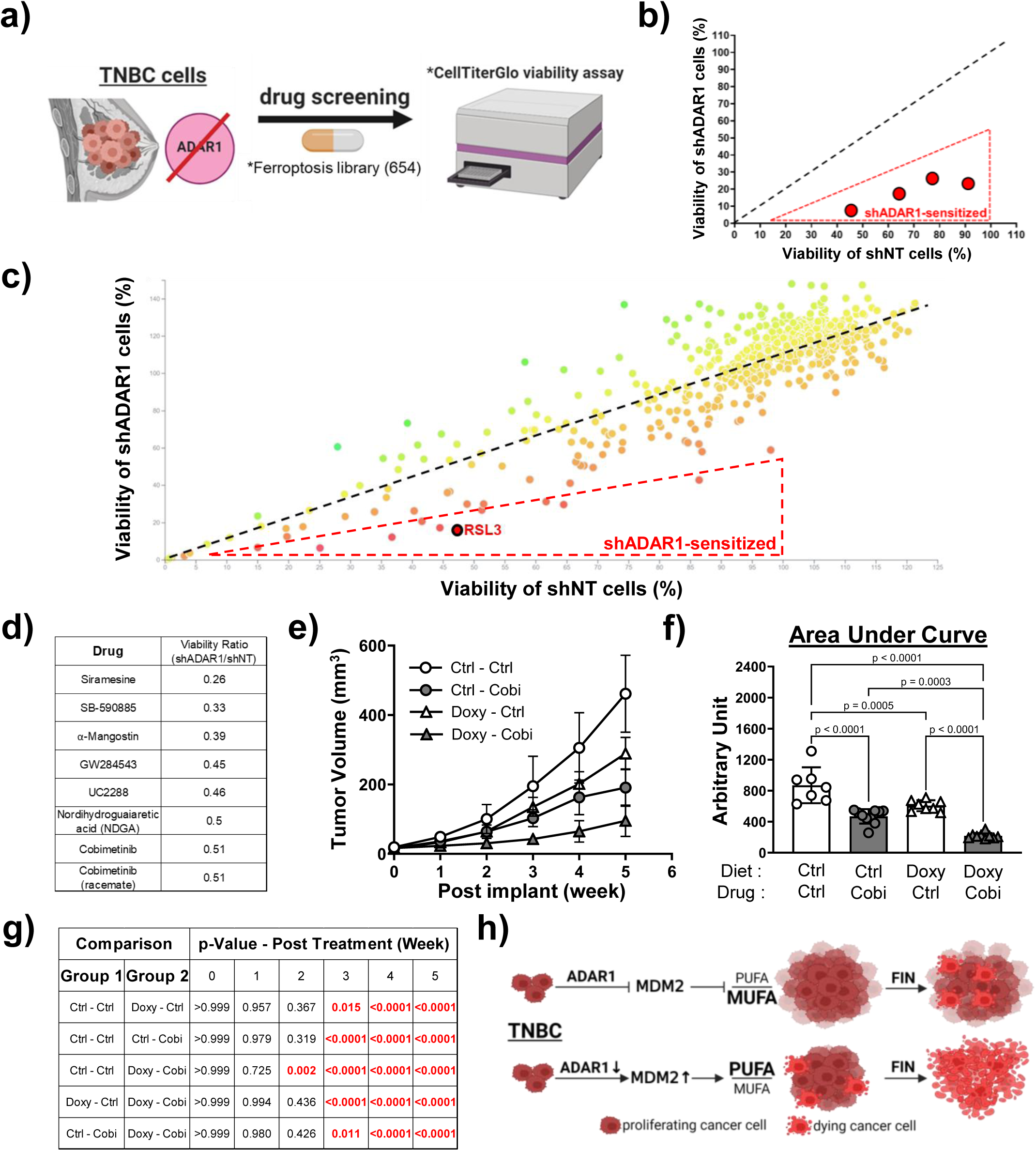
ADAR1 reduction synergizes with cobimetinib to suppress tumorigenesis **A)** Drug screen to identify shADAR1-sensitized ferroptosis modulators. **B)** shADAR1-sensitized hits (red dots) preferentially reduce viability of ADAR1-deficient cells. X and Y axis represent cell viability for ADAR1-intact and -deficient cells, respectively. **C)** Screening result with a ferroptosis-focused library (654 small molecules) in MDA-MB-231 cells. Data were normalized to DMSO-treated cells. RSL3 was included as positive control. Black dashed line denotes expected locations for molecules without discriminating activity. **D)** List of hits (viability ratio shADAR1/shNT ≤ 0.51) identified, ranked by cell viability ratio. **E)** Progression of overall tumor volumes, n=5-10 each group. Error bars mark standard deviation. **F)** Quantification of areas under curve (AUC) from F). Error bars mark standard deviation. **G)** Two-way ANOVA analysis to compare weekly measured overall tumor volumes. Statistically significant p-values (<0.05) were highlighted in red. **H)** ADAR1 protects TNBC cells from ferroptosis through regulating MDM2 and lipid compositions between MUFA and PUFA. FIN, ferroptosis inducer.

Several hits were identified to react at least twice as strongly with ADAR1-deficient cells, compared to ADAR1-intact cells (**Figure 6D**). Siramesine is a lysosome disrupting agent that induces ferroptosis by promoting ROS-mediated oxidative stress and reducing the expression of ferroportin-1, an iron transport protein^33,34^. SB590885 is a B-Raf inhibitor that induces the accumulation of ROS^35^. UC2288 is a derivative of Sorafenib, known to promote ferroptosis by regulating p21 activity^36^. Alpha-Mangostin is a dietary xanthone that induces ROS activity and suppresses lipogenesis^37^. Nordihydroguaiaretic acid (NDGA) is an inhibitor of 5-lipoxygenase (5-LOX) that promotes PUFA peroxidation^38^. GW284543 and cobimetinib are MEK inhibitors, and MEK inhibition has been shown to induce ferroptosis in multiple cancers^39,40^.

Among the identified hits, cobimetinib is the only FDA-approved drug. Cobimetinib is approved for the treatment of metastatic melanoma and histiocytic neoplasms but was also shown to improve the survival of TNBC patients when combined with chemotherapy and immunotherapy^41–43^. Publicly available drug response data show that cobimetinib exhibits stronger activity towards TNBC compared to non-TNBC cell lines, similar to carboplatin, which is known to improve the outcome of TNBC^44^ (**Figure S6A-B**).

The orthotopic implantation mouse model described earlier was used to test the synergistic effect between ADAR1 reduction and cobimetinib *in vivo* (**Figure S6C**). Upon the formation of palpable tumors from MDA-MB-231 cells, mice were fed a control diet or a diet containing doxycycline to induce ADAR1 knockdown before being subjected to cobimetinib treatment (oral gavage at 20mg/kg weight; twice a week) for 5 weeks. The combination of the doxycycline diet and cobimetinib showed superior tumor suppressive efficacy and a stronger ferroptosis phenotype without observed toxicity, indicating a strong synergy between ADAR1 reduction and cobimetinib to suppress tumor progression (**Figure 6E-G, Figure S6D-G**).

## Discussion

Justified by the data shown in this study, we expanded our knowledge of ADAR1-associated vulnerabilities in TNBC by showing that ADAR1 depletion sensitizes TNBC cells to ferroptosis induction. Two recent studies showed that ADAR1 loss leads to overt ferroptosis phenotypes^45,46^. Different experimental conditions are known to affect ferroptosis phenotypes^47^. In particular, CRISPR (clustered regularly interspaced short palindromic repeats)-mediated gene editing used to knock out ADAR1 in these studies has been shown to alter the p53 signaling pathway, which can promote ferroptosis^48,49^. Despite the disparate observations, our results agree with the premise that ADAR1 plays a protective role against ferroptosis.

ADAR1-mediated desensitization to ferroptosis was attributed to lipid remodeling, as the reduction of ADAR1 led to an increased abundance of long and highly unsaturated PUFA, known to be the main drivers of ferroptosis^50^. Transcriptomic analysis revealed that MDM2 is one of the commonly regulated genes in ADAR1-deficient TNBC cell lines that are sensitized to ferroptosis. Consistent with its reported function in promoting ferroptosis through lipid reprogramming, MDM2 was shown to be related to ADAR1-regulated ferroptosis sensitivity^27^. These results reveal a novel mechanism by which ADAR1 protects TNBC cells from stresses that could lead to ferroptosis (**Figure 6H**). Publicly available drug response data show that the sensitivity of TNBC cells to the three ferroptosis inducers used in this study (RSL3, ML162, cobimetinib) is inversely correlated with their dependence on ADAR1 (**Figure S6H-J**). This suggests that the dependence of TNBC cells on ADAR1 could be partially attributed to its ferroptosis-protecting activity, since ferroptosis-tolerant cell lines rely on ADAR1’s protection against ferroptosis induction.

Synthetic lethality, defined as cancer cells acquiring a therapeutically targetable dependency on one gene or pathway for survival following the loss or inhibition of another gene or pathway, has been applied to genetically regulated metabolic vulnerabilities in TNBC^51^. Although recent studies have questioned the status of ADAR1 as a *bona fide* oncogene, it is a promising target for synthetic lethality applications with its ability to suppress multiple stresses directly and indirectly^52^. Our model where ADAR1 inhibition primes TNBC cells for ferroptosis inducers through lipid remodeling presents a novel therapeutic strategy based on recently discovered TNBC biology and ferroptosis-related pharmacology. MUFA has been shown to promote cell proliferation and obesity-driven metastasis of TNBC^53^. Moreover, multiple studies revealed that in TNBC, MUFAs generated intracellularly and by their originating environment protect cancer cells from ferroptosis^54–56^. However, PUFA abundance is positively correlated with ferroptosis sensitivity in TNBC cell lines, and the increase in the PUFA/MUFA ratio has been shown to be effective in sensitizing TNBC to ferroptosis^9,10,57^. A recent study identified an elevated MUFA/PUFA ratio in 3D versus 2D cancer cell cultures, including TNBC, as a major obstacle to translating ferroptosis inducers into preclinical/clinical settings^58^. Our *in vitro* and *in vivo* results shown in this study suggest that ADAR1 inhibition is a promising way to mitigate this challenge.

Preliminary results showed that non-TNBC cells with ADAR1 knockdown are not sensitized to RSL3. Multiple mechanisms contribute to ferroptosis resistance in non-TNBC cells, suggesting higher thresholds for sensitization^59–61^. We posit that the prevalence of p53 mutations in TNBC contributes to this discrepancy, as the ability of MDM2 to promote lipid remodeling and ferroptosis sensitivity could be dictated by its dynamic relationship with p53. The recent discovery that mutant p53 protects TNBC from ferroptosis indicates that MDM2 plays an antagonistic role in promoting ferroptosis sensitivity^62^. With its many p53-independent functions, both pro- and anti-tumorigenic, the complex role of MDM2 in the ADAR1 targeting strategy against TNBC warrants further investigation^63^.

Exciting opportunities are highlighted by the synergistic effect between ADAR1 reduction and ferroptosis induction. In addition to the lack of clinically proven ADAR1 and ferroptosis modulators, both ADAR1 inactivation and ferroptosis have been associated with pathological consequences^32,64,65^. In addition to developing effective ADAR1 inhibitors, functional drug repurposing offers a derisking strategy^66^. A ferroptosis-focused screen identified cobimetinib, an FDA-approved MEK inhibitor with ferroptosis-inducing activity, to be sensitized with ADAR1 reduction. Cobimetinib and other MEK inhibitors have been shown in preclinical and clinical settings to inhibit TNBC tumorigenesis^43,67^. Cobimetinib has also been shown to induce a type I interferon response, indicating the potential of repurposing drugs that affect multiple pathways downstream of ADAR1^68^. Future studies could examine whether other existing TNBC treatments also synergize with ADAR1 inhibition with their ferroptosis-promoting activity.

Recent advances in ADAR1- and TNBC-related research, combined with the results shown in our study, raise further questions about the implication of ADAR1’s ability to regulate lipid metabolism and ferroptosis sensitivity. Does ADAR1 expression or its editing profile have a prognostic value to predict the response of TNBC patients to ferroptosis inducers? As both ferroptosis and ADAR1 inhibition can sensitize ‘immune cold’ tumors to immunotherapy, can a sequential or combined approach targeting ADAR1, ferroptosis, and immune checkpoints minimize resistance to treatment^17,69^? Does ADAR1 protect TNBC from other metabolic stresses, such as amino acid dysregulation? And how do ADAR1-regulated metabolic adaptations fit into dietary interventions to improve cancer treatment^70^? ADAR1 can alter most RNA species and has enormous potential for exploitation to develop innovative therapies against challenging cancers such as TNBC.

## Methods

### Cell culture

Breast cancer cell lines (MDA-MB-231, MDA-MB-436, MDA-MB-468, HCC1806, BT549, BT20, MCF7, T47D, ZR751, SKBR3, BT474, MDA-MB-361) and human embryonic kidney 293T (HEK293T) cells were obtained from American Tissue Cells Consortium (ATCC, Manassas, VA USA). WHIM12 cell line, as part of the patient-derived xenograft (PDX) collection referred to as “Washington University Human in Mouse” (WHIM) lines, was obtained from the Human and Mouse Linked Evaluation of Tumors (HAMLET) core at Washington University in St. Louis. All cell lines were maintained in Dulbecco’s Modification of Eagle’s Medium (DMEM, GE Life Sciences, Pittsburgh, PA USA) supplemented with 10% fetal bovine serum (FBS, Gibco, 10091-148), Sodium Pyruvate (Cellgro, Waltham, MA USA, 30-002-CI), Non-Essential Amino Acids (NEA, Cellgro, 25-030-CI), and L-glutamine (Cellgro, 25-005-CI). Sensitivity of cells to ferroptosis inducing agents were tested in defined medium with low serum using DMEM supplemented with 2% FBS and L-glutamine. LipoFexin (Lamda Biotech, St. Louis, MO USA, TS310) was used to generate lentivirus. Fugene 6 transfection reagent (Promega, E2692) was used for other transfection experiments.

### Plasmids

Tetracycline-inducible shADAR1 was generated by cloning shADAR1 oligonucleotides (targets 3’UTR: GCCCACTGTTATCTTCACTTT) into Tet-pLKO.1-puro construct (Addgene #21915) with AgeI and EcoRI restriction enzyme sites. Tet-inducible MDM2-overexpressing construct was generated by subcloning MDM2 open-reading frame (ORF) from pCMV-myc3-MDM2 (Addgene #20935) into pSBtet-GP sleeping beauty vector (Addgene #60495) using the Sfil restriction enzyme site. pSBtet-GP-MDM2 and pCMV(CAT)T7-SB100 transposase construct (Addgene #34879) were co-transfected into MDA-MB-231 cells to generate a puromycin-resistant stable cell line. Constructs containing shADAR1 (pLKO.1-shADAR1) and shNT (pLKO.1-shNT; CCTAAGGTTAAGTCGCCCTCG) were obtained from MilliporeSigma (MISSION^®^ Lentiviral shRNA; TRCN0000050788) and Addgene (#136035), respectively.

### Lentiviral production and transduction

To generate lentivirus, HEK293T cells were transfected using LipoFexin with pCMV-VSV-G (Addgene #8454), pCMV-ΔR8.2 (Addgene #8455), and expression constructs. Growth medium was replaced with fresh medium 24hr after transfection, and supernatants containing lentivirus were harvested 24hr later. For transduction, one million cells were infected with lentivirus for 24hr in the presence of 10 µg/ml protamine sulfate to facilitate viral entry. Transduced cells were selected with puromycin (2 µg/ml) (Gibco, A11138-03).

### Cell viability assay

For cell viability assays, cells were plated at a density of 1,000 to 3,000 cells per well on a 96-well plate and left to grow overnight in complete growth medium at 37°C. Cells were then treated with the indicated drugs in equal volume at 2× desired concentration and left to incubate for 72 h. Cells were then incubated with CellTiter-Glo® 2.0 Reagent (Promega, Madison, WI, USA, G924C) for 1h at room temperature, and viability was read out according to the manufacturer’s protocol on a GloMax Navigator Microplate Luminometer (Promega, GM2000). The half-maximal inhibitory concentration (IC_50_) was determined using a nonlinear regression model using Prism 10 software (GraphPad, https://www.graphpad.com).

Chemicals used during viability assays in this study include RSL3 (Selleck chemicals, #S8155), ML162 (Selleck chemicals, #S4452), UAMC-3203 (MedChemExpress, #HY-112909), Ferrostatin-1 (Selleck chemicals, #S7243), Liproxstatin-1 (Selleck chemicals, #S7699), SP-141 (MedChemExpress, #HY-110182), MEL23 (MilliporeSigma, #373227), MX69 (Selleck chemicals, #S8403), and pirinixic acid (MedChemExpress, #HY-16995).

### Lipid peroxidation assay; Glutathione activity assay; Ferrous iron assay

Biochemical assays were performed to evaluate ferroptosis phenotypes or molecular mechanisms contributing to cells’ sensitivity to ferroptosis induction. Level of lipid peroxidation was determined by **1)** measuring its natural byproduct malondialdehyde (MDA) using MDA Colorimetric/Fluorometric Assay (Abcam, Cambridge, UK, ab118970), or by **2)** staining and comparing levels of reduced and oxidized forms of lipid peroxidation probe BODIPY^TM^ 581/591 C11 (Image-iT^TM^ Lipid Peroxidation Kit, Thermo Fisher Scientific, #C10445), according to manufacturer’s instruction. Intracellular oxidative stress level was determined by measuring the ratio between reduced (GSH) and oxidized (GSSG) forms of Glutathione using GSH/GSSG Ratio Detection Assay (Abcam, ab205811) according to manufacturer’s instruction. Ferrous iron, which drives ferroptosis, was measured by FerroOrange staining (Dojindo, Japan, F374) according to the manufacturer’s instructions. Quantification of signal intensity was performed using ImageJ software (NIH, Bethesda, MD USA).

### Untargeted lipidomic analysis with reversed-phase liquid chromatography (RPLC)

Cells were washed with cold PBS 3 times, cold molecular biology grade water (Corning, Glendale, AZ, 46-000-CM) 3 times, metabolically quenched with cold methanol for 5 min, scraped harvested, and centrifuged at 10,000 × g for 5 min to remove supernatant. Cell pellets were dried using CentriVap Concentrator (Labconco, Kansas City, MO, 79700-00) and stored in −80°C before extraction of metabolites. Cold extraction buffer containing Methanol:Acetonitrile:Water at 2:2:1 ratio was added to the pellets. Samples were then subjected to 3 times of 30-sec vortex → 1-min liquid nitrogen bath → 10-sec thaw → 10-min bath sonication at 25°C (Fischer Scientific, CPXH Series Model 3800 Heated Ultrasonic Cleaner), before being stored at −20°C for 2 hours to overnight. After centrifuging at 14000 RPM at 4°C for 10 min, sample supernatants were transferred to new Eppendorf tubes and speed dried for 2 hours. BCA Protein Assay was performed on cell pellets in advance to determine protein concentration. Isopropanol:Methanol at 1:1 ratio was added at 1µL per 2.5µg of proteins to the residue and subjected to 2 times of 5-min bath sonication at 25°C → 1-min vortex, before being stored at 4°C for 1 hour to overnight. After centrifuging at 14000 RPM at 4°C for 10 min, supernatants were transferred to LC vials for LC/MS analysis. Following extraction, an internal standard (Avanti Research, SPLASH® LIPIDOMIX® Mass Spec Standard, 330707) was spiked into each experimental sample for quality control.

An aliquot (4µL) of the extracted sample was injected onto a Waters HSS T3 1.8µM, 2.1×150mm column with a 1.8µM, 2.1×50mm guard column connected to 1290 Infinity II LC. The columns were used with the following buffers and gradients: Mobile Phase A = 40% water, 60% Acetonitrile, 0.1% Formic Acid, 10mM Ammonium Formate, 2.5µM Medronic Acid; Mobile Phase B = 90% Isopropanol, 10% Acetonitrile, 0.1% Formic Acid, 10mM Ammonium Formate; Needle Wash = 70% Mobile Phase A solution + 30% Mobile Phase B solution; Seal Wash = 9:1 Water:Isopropanol. All solutions were mixed and degassed for 10 min before use. The binary pump was operated at 70% A + 30% B from 0 - 17 min, 25% A + 75% B 0% B from 17 - 20 min, 15% A + 85% B from 20 - 23 min, 100% B from 23 - 27 min, and 70% A + 30% B thereafter at constant 0.25 mL/min flow rate. MS detection was carried out using Agilent 6545 Q-TOF with 150-1,200 Da mass range. Lipid Annotator (Version 1.0, Agilent) was used to generate a target list for all lipids detected. LC/MS results were analyzed and visualized with MetaboAnalyst (McGill University, https://www.metaboanalyst.ca).

### Bulk RNA sequencing and analysis

Total RNA was isolated from cells transduced with shNT or shADAR1 lentivirus for 24hr, followed by 24hr of puromycin selection, using miRNeasy Mini Kit (Qiagen, Hilden, Germany, 217004) including on-column DNase digestion following the manufacturer’s protocol. Samples were prepared according to RiboErase library kit (Roche) manufacturer’s protocol, indexed, pooled, and sequenced on an Illumina NovaSeq S4. Base calls and demultiplexing were performed with Illumina’s bcl2fastq software and a custom python demultiplexing program with a maximum of one mismatch in the indexing read. RNA-seq reads were then aligned to the Ensembl release 76 primary assembly with STAR version 2.5.1a1. Gene counts were derived from the number of uniquely aligned unambiguous reads by Subread:featureCount version 1.4.6-p52. Sequencing performance was assessed for the total number of aligned reads, total number of uniquely aligned reads, and features detected. All gene counts were then imported into the R/Bioconductor package EdgeR and TMM normalization size factors were calculated to adjust for samples for differences in library size. Ribosomal genes and genes not expressed in the smallest group size minus one sample greater than one count-per-million were excluded from further analysis. The TMM size factors and the matrix of counts were then imported into the R/Bioconductor package Limma. Weighted likelihoods based on the observed mean-variance relationship of every gene and sample were then calculated for all samples with voomWithQualityWeights. The performance of all genes was assessed with plots of the residual standard deviation of every gene to their average log-count with a robustly fitted trend line of the residuals. Differential expression analysis was performed to analyze differences between conditions and raw p-values and Benjamini-Hochberg false-discovery rate (FDR) adjusted p-values were determined to filter results based on cut-off. Biological and functional pathways analyses, as well as gene set enrichment analyses were performed with the Enrichr platform (Icahn School of Medicine at Mount Sinai, https://amp.pharm.mssm.edu/Enrichr), the GeneOntology enrichment tools (Gene Ontology Consortium, https://geneontology.org/docs/go-consortium), and the Gene Set Enrichment Analysis (GSEA) software (Broad Institute/UC San Diego, https://www.gsea-msigdb.org/gsea). p<0.05 was used as the cutoff to identify genes commonly regulated among 4 TNBC cell lines, and FDR<0.1 was used as the cutoff to identify regulated genes in MDA-MB-231 cells for GSEA analysis. Relevant molecular signatures databases (MSigDB) were selected for GSEA analysis. When multiple MSigDB files exist for a similar pathway, the one with the biggest gene list was preferred. MSigDB used in this study include WP_FERROPTOSIS, REACTOME_FATTY_ACID_METABOLISM, HALLMARK_REACTIVE_OXYGEN_SPECIES_PATHWAY, REACTOME_IRON_UPTAKE_AND_TRANSPORT, REACTOME_CELL_CYCLE, REACTOME_INTERFERON_SIGNALING, HALLMARK_GLYCOLYSIS, and GOBP_AMINO_ACID_METABOLIC_PROCESS.

### Immunoblotting (western blotting) analysis

Cells were washed with phosphate-buffered saline (PBS, Corning, #21-031-CM), scrape harvested, centrifuged at 1000 × g for 5 min, and lysed with RIPA buffer (20mM Tris-HCl pH7.5, 150mM NaCl, 1mM EDTA, 1% NP-40, 0.1% SDS, 0.1% Deoxycholate) supplemented with 10mM PMSF and HALT protease inhibitor cocktail (Thermo Fisher Scientific, #78440). Lysates were clarified by centrifugation, and the protein concentration was determined using DC protein assay system (Bio-Rad, Hercules, CA USA, #5000113). Equal amount of protein was resolved by sodium dodecyl sulfate-polyacrylamide gel electrophoresis (SDS-PAGE) using Criterion TGX Stain-Free Precast Gels (Bio-Rad, #5678094) and transferred onto Immobilon-P membranes (MilliporeSigma, ISEQ85R). Primary antibodies used in this study include anti-ADAR1 antibody (Santa Cruz Biotechnology, sc-73408; RRID:AB_2222767; 1:300), anti-GAPDH antibody (Bethyl Laboratories, A300-641A; RRID:AB_513619; 1:10,000), anti-MDM2 antibody (Cell Signaling Technology, #86934; RRID:AB_2784534; 1:1,000), and anti-Myc-tag antibody (Cell Signaling Technology, #2278; RRID:AB_490778; 1:1,000). Secondary antibodies conjugated to Horseradish peroxidase were used at a dilution of 1:5-10,000 (Jackson Immunochemicals, West Grove, PA USA). Clarity Western ECL Substrate (Bio-Rad, #170-5061) was then applied to blots and protein levels were detected using autoradiography with ChemiDoc XRS+ Imager (Bio-Rad).

### RNA editing site-specific quantitative PCR (RESS-qPCR)

Total RNA was isolated from lentivirus-transduced cells using RNeasy Plus Mini Kit including on-column DNase digestion following the manufacturer’s protocol. To eliminate any residual genomic DNA contamination, TURBO DNA-free^TM^ Kit (ThermoFisher Scientific, AM1907) was used to further clean up isolated RNAs. RNAs free of genomic DNA were reverse transcribed to cDNAs using the High-Capacity cDNA Reverse Transcription Kit (ThermoFisher Scientific, #4368814). Primers were designed to amplify MDM2-WT (Forward: ATAGGACTGAGGTAATTCTGCACAGCA; Reverse: ATAATGCTTGGAGGACCTCCACATGT) and MDM2-Edited (Forward: TAAATGGCCAAAGGGATTAGTAGTGTG; Reverse: AAGAGATTCTGCTTGGTTGTAGCTGAAG). Quantitative PCR was performed using iTaq Universal SYBR Green Supermix (Bio-Rad, #1725125) on the CFX96 Touch Real-Time PCR Detection System (Bio-Rad, #1845097). The relative RNA editing level (Edit/Wild-Type RNA ratio) was calculated as 2^-(Ct^ ^Edit^ – ^Ct^ ^WT)^.

### Sanger sequencing to detect RNA editing

Isolated RNAs free of genomic DNA were used for Sanger sequencing analysis to verify RNA editing events. PCR was performed using the high-fidelity Phusion PCR kit according to the manufacturer’s instructions (New England Biolabs, Ipswich, MA, E0553). “Outer” primers were used to produce amplicons flanking the editing site about 50-150 bp on either side to ensure successful sequencing analysis: MDM2 3’UTR Motif 1 (Forward: GCCTAGCAATGATCTAGAAGCAG; Reverse: GCACACCCATCAGCTCAAATAT), MDM2 3’UTR Motif 2 (Forward: ATAGGACTGAGGTAATTCTGCACAGCA; Reverse: AAGAGATTCTGCTTGGTTGTAGCTGAAG). Successful PCRs were verified by DNA gel electrophoresis and amplicons were extracted from the gel using Monarch gel extraction kit (BioRad, T1020S) according to manufacturer’s instructions and submitted for sequencing analysis (IDT Technologies). Sanger sequencing was performed using the outer primers (A/G were read for WT/Edited if forward primers were used; T/C were read if reverse primers were used) used for PCR, and sequence chromatograms were analyzed and converted to A/G reading for presentation purposes using Chromas software (Technelysium, https://technelysium.com.au/wp/chromas).

### Drug screening with ferroptosis-focused library

MDA-MB-231 cells treated with shNT (ADAR1-intact) or shADAR1 (ADAR1-deficient) were screened against the ferroptosis-focused drug library (MedChemExpress, HY-L051) at the High Throughput Screening Center (HTSC) at Washington University in St. Louis. CellTiter-Glo® Viability Assay was utilized to determine cell viability upon drug treatment. Cells were prepared in defined medium with low serum using DMEM supplemented with 2% FBS and L-glutamine.

All drugs in the library were pre-dispensed into 96-well sterile polypropylene plates (Corning, Axyen) using a Hummingbird and subsequently diluted with media to produce intermediate plates at 20µM and 2µM. Diluted compounds were then transferred using a BiomekFX (Beckman Coulter) to 96-well white cell culture plates (Greiner Bio-One, Monroe, NC, 655083) (50µl/well, 18 plates total with 9 plates for each experimental group. Select wells containing DMSO alone served as controls. Cells were seeded (15,000 cells in 50µl cell culture media per well) using the Multidrop 384 dispenser (Thermo Fisher Scientific, 24291). Plates were kept at room temperature for 30 min before being moved to the incubator (37°C, 5% CO_2_) for 24h. On day 2, viability Assays were performed using the integrated and automated platform (Beckman Coulter), with library plate-matched ADAR1-intact and ADAR1-deficient samples alternating between screening steps: Cell plates were equilibrated at room temperature for 30 min in a Cytomat MPH incubator. CellTiter-Glo® reagent (100µl/well) was added with a Multidrop 384 dispenser. Plates were shaken at 250 rpm for 2 min with a Teleshake followed by room temperature incubation for 1h in a Cytomat MPH incubator. Luminescence was measured (1 sec/well after a 1 min delay) with a Perkin Elmer Envision plate reader. To monitor assay performance, Coefficient of variations (CVs) and Z’-factors for DMSO-treated and no-cell control samples were determined. All screening plates were confirmed to display high quality assay metrics (CVs < 10%; Z’-factors > 0.8). All test wells were normalized to no-treatment samples on the same plate. Relative cell viability was compared between ADAR1-intact and ADAR1-deficient samples, and the cut-off was set at 51% to discriminate hits from no-hits. Drug screening results were analyzed and visualized with CDD vault (Collaborative Drug Discovery, https://www.collaborativedrug.com).

### Mammary fat pad orthotopic implantation

Immuno-deficient NOD scid gamma (NSG) female mice at 6-8 weeks old were purchased from Jackson Laboratory (Bar Harbor, ME, USA, #005557) and placed on Teklad Global 18% protein rodent diet (Envigo, Indianapolis, USA, T.2918.15) 3 days before implantation. 1.5 x 10^5^ MDA-MB-231 cells were harvested and resuspended in PBS, mixed with Matrigel® Matrix Basement Membrane (Corning, Waltham, USA, #354230) at 1:1 volume ratio, and kept at 4°C until implantation. In total 100µl of cells-Matrigel solutions were injected into the right inguinal mammary glands of NSG mice. Ten days after the implantation, palpable tumors were detected and half of the mice were switched to doxycycline-containing diet (Envigo, 625mg/kg, TD.01306). RSL3 (100mg/kg weight; IP injection) and cobimetinib (MedChemExpress, HY-13064; 20mg/kg weight; Oral gavage) were freshly prepared in DMSO:Corn oil (MedChemExpress, HY-Y1888) before treatment, starting 3 days after the diet switch. Tumors were measured using a caliper once per week, and tumor volumes were calculated using the following formula: L × W^2^ × 0.52 (L and W represent the large and small diameter of the tumor, respectively). Mice were euthanized 6 weeks post implantation or before tumors reached 2 cubic cm in size. Tumors were dissected from the mice for weight measurement and subsequently fixed in 10% formalin, before transferred to 70% ethanol for processing to prepare formalin-fixed paraffin-embedded (FFPE) tissue blocks. All animal-related experimental procedures were performed in compliance with the guidelines given by the American Association for Accreditation for Laboratory Animal Care and the U.S. Public Health Service Policy on Human Care and Use of Laboratory Animals. All animal studies were approved by the Washington University Institutional Animal Care and Use Committee (IACUC) in accordance with the Animal Welfare Act and NIH guidelines (Protocol 20220144).

### Immunohistochemistry (IHC)

Tumor FFPE tissue blocks were sectioned at 5μm thickness. Hematoxylin and eosin (H&E) staining was performed for histological evaluations. For immunohistochemistry, sections were stained using a Bond RXm autostainer (Leica, Buffalo Grove, IL USA). Primary antibodies for ADAR1-p150 (Abcam, ab126745; RRID:AB_11145661) and 4-Hydroxynonenal (4HNE, ThermoFisher Scientific, MA5-27570; RRID:AB_2735095) were diluted 1:100 in Antibody Diluent (Leica, AR9352). Briefly, slides were baked at 65°C for 15min and automated software performed dewaxing, rehydration, antigen retrieval, blocking, primary antibody incubation, post primary antibody incubation, detection (DAB chromogen; 3,3′-diaminobenzidine), and hematoxylin counterstaining using Bond reagents (Leica, DS9800 system). Samples were then removed from the machine, dehydrated through ethanol and xylenes, mounted and cover slipped before being imaged using Zeiss Axioskop 2 MOT microscope. Quantification of DAB signals was performed using ImageJ Fiji package.

### Bioinformatic analysis and data availability

Bioinformatic analyses to assess relationships among gene expression, drug response, gene dependency and metabolomics in breast cancer cell lines were performed using the DepMap public data (depmap.org) with the TNBC (HCC1187, HS578T, HCC1599, DU4475, BT549, HCC1143, BT20, MDA-MB-436, HCC1806, HCC70, HCC2157, HCC1395, MDA-MB-231, MDA-MB-468, MDA-MB-453) and non-TNBC (SKBR3, MCF7, KPL1, MDA-MB-134VI, ZR751, EFM192A, T47D, AU565, HCC1419, EFM19, HCC1500, HCC1428, UACC893, UACC812, HCC202, HCC2218, MDA-MB-175VII, BT483, ZR7530, MDA-MB-415, BT474, HCC1569, MDA-MB-361) groups. The results published here are in whole or part based upon data generated by Cancer Target Discovery and Development (CTD²) Network (https://ccg.cancer.gov/ccg/research/functional-genomics/ctd2/data-portal) established by the National Cancer Institute’s Center for Cancer Genomics. Kaplan-Meier plots for breast cancer patients’ survival outcome were obtained from Kaplan-Meier Plotter (https://KMplot.com), using the mRNA gene chip data with auto select best cutoff. The secondary RNA structure of MDM2 3’UTR was predicted and visualized with RNAfold web server (ViennaRNA Web Services, https://rna.tbi.univie.ac.at). RNA-seq data from this study are available at the Gene Expression Omnibus under accession number GSE286185.

## Statistical analysis

All *in vitro* and *in vivo* data are reported as the mean ± SD unless stated otherwise, and statistical analyses were performed using GraphPad Prism 10. The two-tailed unpaired student t test was performed for experiments comparing between two experimental groups. One-way ANOVA with Tukey’s multiple comparisons was used for statistical analysis of all other experiments unless stated otherwise. Tumor volume analyses were performed by **1)** comparing AUC (area under curve) between experimental groups using one-way ANOVA, and **2)** comparing weekly measured tumor volumes between groups using two-way ANOVA with Tukey’s multiple comparison. Pearson correlation coefficient and/or Spearman’s rank correlation coefficient were calculated to assess correlations between two data sets. Statistical significance was evaluated using p-value, which is shown in the numerical format. Lower p-values reject the null hypothesis more strongly, and p-value <0.05 is generally considered statistically significant.

## Resource availability

### Lead contact

Requests for further information and resources should be directed to and will be fulfilled by the lead contact, Jason D. Weber.

### Materials availability

All unique/stable reagents generated in this study are available from the lead contact with a completed materials transfer agreement.

### Data and code availability

All data reported in this paper will be shared by the lead contact upon request. This paper does not report original code. Any additional information required to reanalyze the data reported in this paper is available from the lead contact upon request.

## Acknowledgements

We thank the Division of Comparative Medicine at Washington University in St. Louis for housing the mouse colony. We thank the Denardo lab for processing the fixed tumor tissues. We thank the Histology and Morphometry Core, Musculoskeletal Research Center (supported by NIH P30 AR074992), for sectioning the tumor samples and performing H&E staining. We thank the support by the Genome Technology Access Center (GTAC) at McDonnell Genome Institute (MGI) Symposium Pilot Project Funding to perform RNA-seq analyses (Catrina Fronick and Michael Heinz for logistics; Toni Sinnwell, Elliott Klotz, and Eric Tycksen for quality control, sequencing and analyses). We acknowledge the Just-In-Time (JIT) Core Usage Funding Program to support the collaboration with HTSC for drug screening. Both GTAC@MGI Symposium Pilot Project and JIT programs are funded through the Institute of Clinical and Translational Sciences (ICTS), Washington University in St. Louis. ICTS was supported by the National Center for Advancing Translational Sciences (NCATS) grant UL1TR002345. This work was supported by R01CA262804 (JDW) from National Cancer Institute (NCI), and W81XWH-21-1-0476 and W81XWH-21-1-0466 from the Congressionally Directed Medical Research Programs (CDMRP), Department of Defense (JDW). C-P. K received support from Transdisciplinary Research in Energetics and Cancer (TREC) Training Workshop R25CA203650 from NCI and TL1 Translational Sciences Postdoctoral Program TL1TR002344 from National Center for Advancing Translational Sciences. L.S.T. is supported by R25HG006687 and T32GM148405 from National Human Genome Research Institute and National Institute of General Medical Sciences, respectively.

## Author contributions

**C-P. Kung:** Conceptualization, investigation, methodology, formal analysis, project administration, writing–original draft, writing–review and editing. **N.D. Terzich:** investigation, methodology, formal analysis. **M.X.G. Ilagan:** investigation, methodology, formal analysis, writing–review and editing. **M.J. Prinsen:** Investigation, methodology, formal analysis. **M. Kaushal:** Investigation, methodology, formal analysis. **R.D. Kladney:** Investigation, methodology. **J.H. Weber:** Investigation. **A.R. Mabry:** Investigation. **L.S. Torres:** Investigation. **E.R. Bramel:** Investigation. **E.C. Freeman:** Investigation. **T. Sabloak:** Investigation. **K.A. Cottrell:** Investigation. **S. Ryu:** Investigation. **W.M. Weber:** Investigation. **L.B. Maggi Jr.:** investigation, writing–review and editing. **L.P. Shriver:** investigation, methodology, writing–review and editing. **G.J. Patti:** funding acquisition, investigation, writing–review and editing. **J.D. Weber:** Conceptualization, resources, supervision, funding acquisition, writing–original draft, writing–review and editing.

## Supplementary information

Supplementary Figure S1-S6

Supplementary Table 1. Genes altered in ADAR1-deficient TNBC cell lines

Supplementary Table 2. Signaling pathways altered upon ADAR1 knockdown

Supplementary Table 3. Disease-related biological functions altered upon ADAR1 knockdown

Supplementary Table 4. Biological pathways altered upon ADAR1 knockdown

