## Supplementary Figure 1-6 for "ADAR1 Regulates Lipid Remodeling to Dictate Ferroptosis Sensitivity"

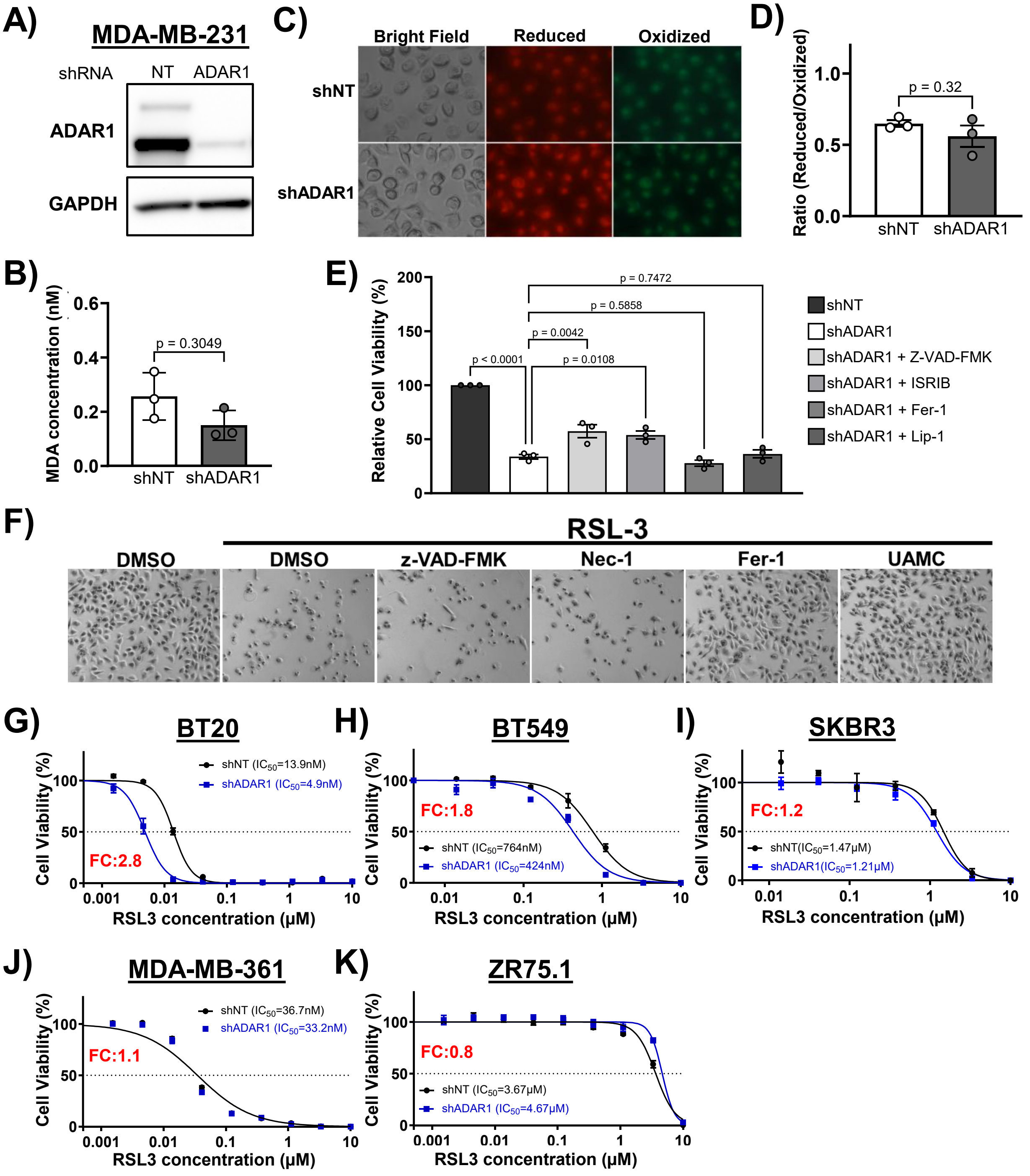
A)
MDA-MB-231
shRNA
NT
ADAR1
ADAR1
GAPDH
B)
MDA concentration (nM)
p = 0.3049
0.0
0.2
0.4
0.6
shNT
shADAR1
C)
Bright Field
Reduced
Oxidized
shNT
shADAR1
D)
Ratio (Reduced/Oxidized)
p = 0.32
0.0
0.5
1.0
shNT
shADAR1
E)
Relative Cell Viability (%)
0
50
100
150
200
p < 0.0001
p = 0.0042
p = 0.0108
p = 0.5858
p = 0.7472
shNT
shADAR1
shADAR1 + Z-VAD-FMK
shADAR1 + ISRIB
shADAR1 + Fer-1
shADAR1 + Lip-1
F)
RSL-3
DMSO
DMSO
z-VAD-FMK
Nec-1
Fer-1
UAMC
G)
BT20
Cell Viability (%)
shNT (IC50=13.9nM)
shADAR1 (IC50=4.9nM)
FC:2.8
0.001
0.01
0.1
1
10
RSL3 concentration (μM)
H)
BT549
Cell Viability (%)
FC:1.8
shNT (IC50=764nM)
shADAR1 (IC50=424nM)
0.01
0.1
1
10
RSL3 concentration (μM)
I)
SKBR3
Cell Viability (%)
FC:1.2
shNT(IC50=1.47μM)
shADAR1(IC50=1.21μM)
0.01
0.1
1
10
RSL3 concentration (μM)
J)
MDA-MB-361
Cell Viability (%)
shNT (IC50=36.7nM)
shADAR1 (IC50=33.2nM)
FC:1.1
0.001
0.01
0.1
1
10
RSL3 concentration (μM)
K)
ZR75.1
Cell Viability (%)
FC:0.8
shNT (IC50=3.67μM)
shADAR1 (IC50=4.67μM)
0.001
0.01
0.1
1
10
RSL3 concentration (μM)

**Figure S1. ADAR1 protects TNBC cells from ferroptosis induction**

**A)** Immunoblot analysis showed significant ADAR1 reduction in MDA-MB-231 cells treated with shADAR1-containing lentivirus. Images are representative of three replicates. GAPDH, loading control. **B)** Concentration of lipid peroxidation marker MDA in MDA-MB-231 cells. n=3. Error bars mark standard deviation. **C)** Light and fluorescent images showed morphology (Bright Field) and reduced and oxidized forms of lipid peroxidation probe BODIPY$^{TM}$ 581/591 C11 in MDA-MB-231 cells. **D)** Quantification of ratios of reduced over oxidized forms of BODIPY$^{TM}$ 581/591 C11. n=3. Error bars mark standard error. **E)** Relative cell viability of ADAR1-intact and -deficient MDA-MB-231 cells, as well as ADAR1-deficient cells treated with apoptosis inhibitor Z-VAD-FMK (50μM), eIF2α inhibitor ISRIB (10μM), and ferroptosis inhibitors Fer-1 (Ferrostatin-1, 1μM) and Lip-1 (Liproxstatin-1, 1μM) for 72h. n=3. Error bars mark standard error. **F)** Light micrographs of MDA-MB-231 cells treated with RSL3 (10μM), in combination with DMSO, Z-VAD-FMK (5μM), necroptosis inhibitor Nec-1 (Necrostatin-1, 5μM), and ferroptosis inhibitors Fer-1 (5μM) and UAMC (5μM) for 24h. $IC_{50}$ values of RSL3 were determined for ADAR1-intact and -deficient **G)** BT20 and **H)** BT549 TNBC cell lines, as well as **I)** SKBR3, **J)** MDA-MB-361, and **K)** ZR75.1 non-TNBC cells. Graphs are representatives of three independent experiments.

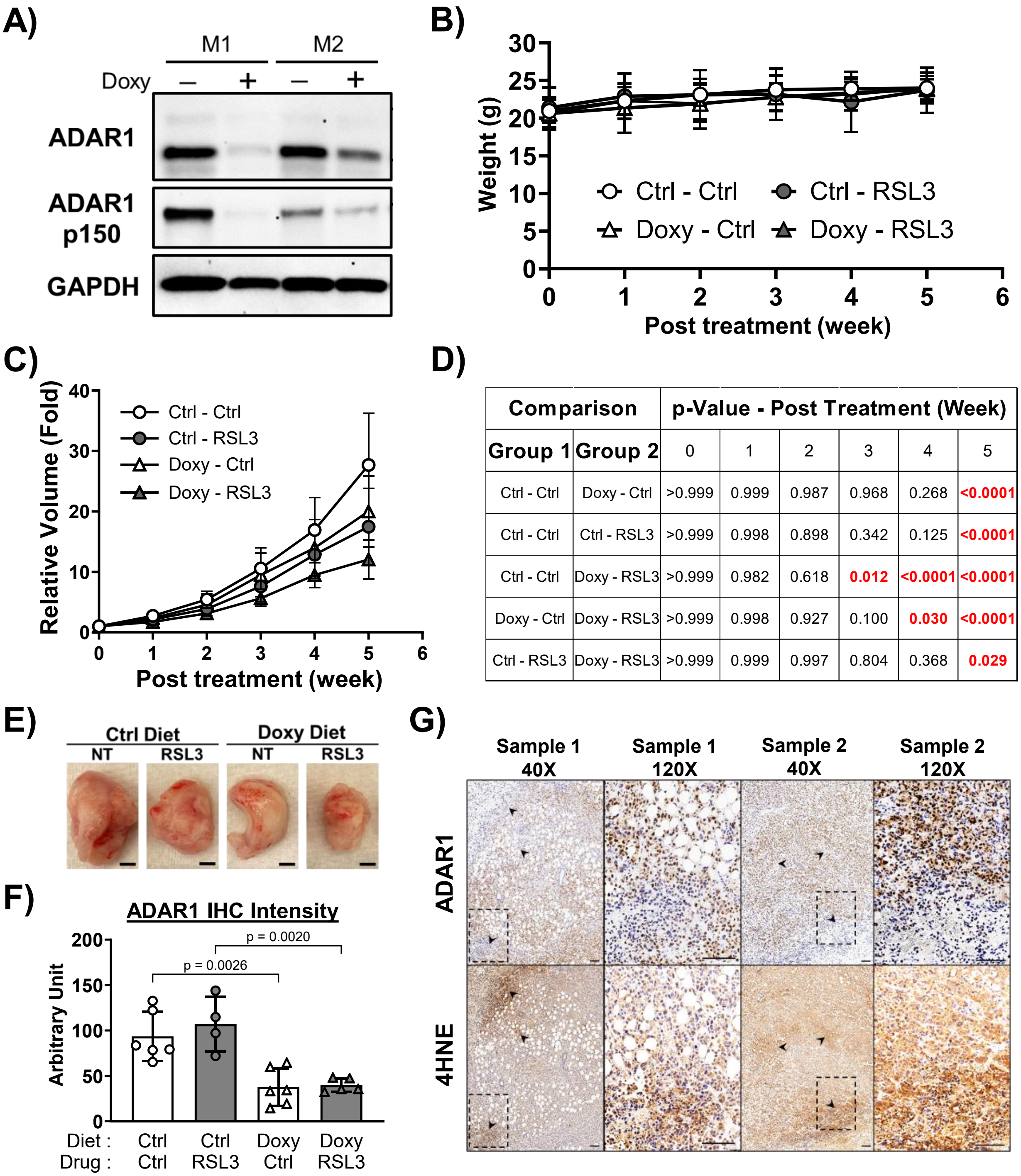


| Comparison | | p-Value - Post Treatment (Week) | | | | | |
|---|---|---|---|---|---|---|---|
| Group 1 | Group 2 | 0 | 1 | 2 | 3 | 4 | 5 |
| Ctrl - Ctrl | Doxy - Ctrl | >0.999 | 0.999 | 0.987 | 0.968 | 0.268 | <0.0001 |
| Ctrl - Ctrl | Ctrl - RSL3 | >0.999 | 0.998 | 0.898 | 0.342 | 0.125 | <0.0001 |
| Ctrl - Ctrl | Doxy - RSL3 | >0.999 | 0.982 | 0.618 | 0.012 | <0.0001 | <0.0001 |
| Doxy - Ctrl | Doxy - RSL3 | >0.999 | 0.998 | 0.927 | 0.100 | 0.030 | <0.0001 |
| Ctrl - RSL3 | Doxy - RSL3 | >0.999 | 0.999 | 0.997 | 0.804 | 0.368 | 0.029 |

**Figure S2. Combining ADAR1 reduction with RSL3 suppresses orthotopically implanted MDA-MB-231 xenograft progression *in vivo***

**A)** Immunoblot analysis showed ADAR1 reduction in lysates extracted from newly formed tumors from MDA-MB-231 cells possessing Tet-inducible shADAR1, 7 days after implanted mice (M1 and M2) were switched to doxycycline diet. GAPDH, loading control. **B)** Progression of average mouse weight. n=5-9 each group. Error bars mark standard deviation. **C)** Progression of relative tumor volumes, normalized to week 0. n=5-9 each group. Error bars mark standard deviation. **D)** Two-way ANOVA analysis compares weekly measured relative tumor volumes. Statistically significant p-values (<0.05) were highlighted in red. **E)** Representative images of tumors resected after the treatment regimens concluded. Scale bar, 5mm. **F)** Quantification of IHC staining intensity of ADAR1. Error bars mark standard deviation. N=4-6 each group. **G)** Representative images of IHC staining of ADAR1 and 4HNE in 2 resected tumor samples from Doxy-RSL3 mice. Black arrows point to areas where staining intensity between ADAR1 and 4HNE are inversely correlated. Black dashed boxes were zoomed in on the right. Scale bar, 200μm.

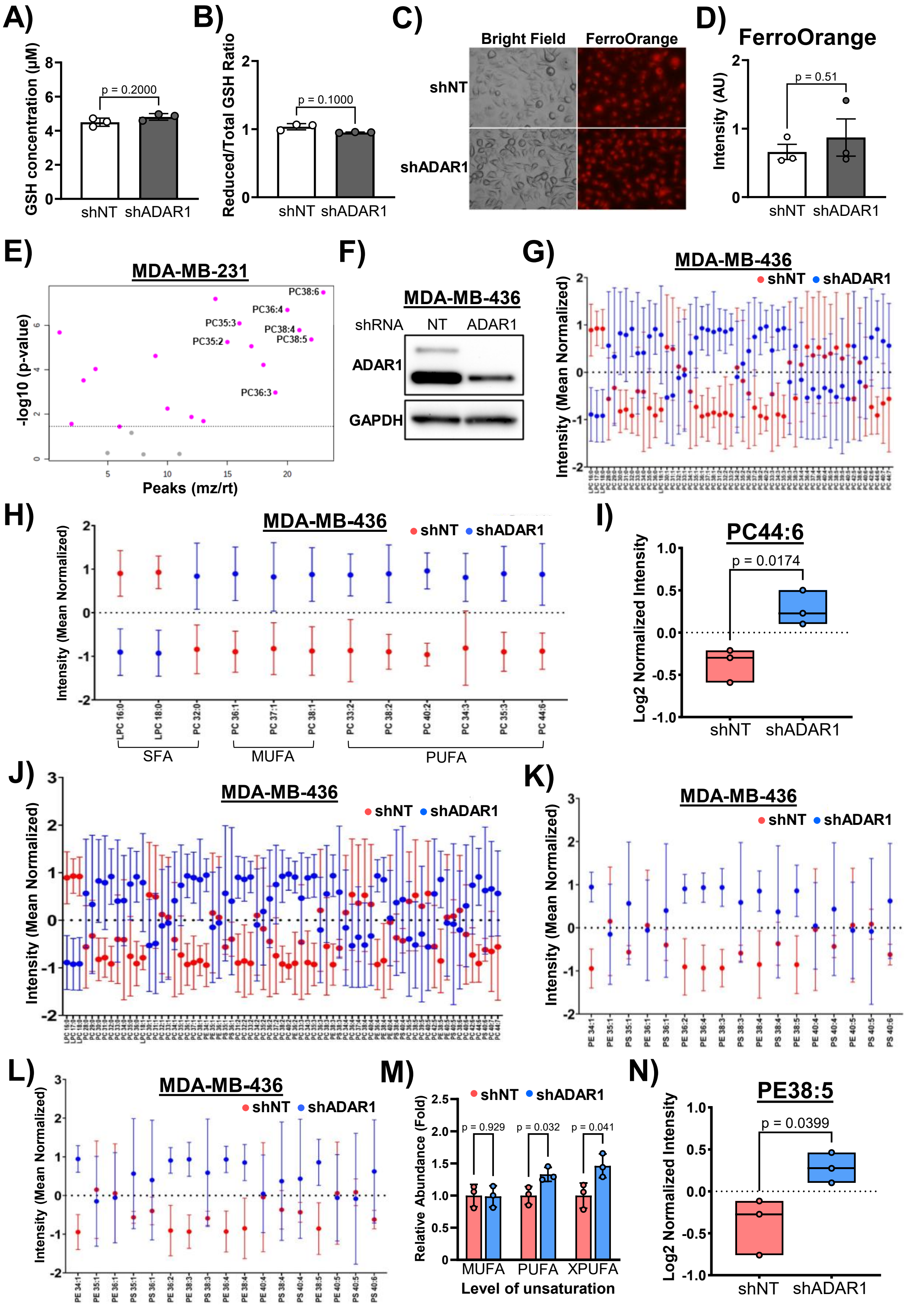
A)
GSH concentration (μM)
p = 0.2000
shNT shADAR1
B)
Reduced/Total GSH Ratio
p = 0.1000
shNT shADAR1
C)
Bright Field
FerroOrange
shNT
shADAR1
D)
FerroOrange
Intensity (AU)
p = 0.51
shNT shADAR1
E)
MDA-MB-231
-log10 (p-value)
Peaks (mz/rt)
PC38:6
PC36:4
PC35:3
PC38:4
PC35:2
PC38:5
PC36:3
F)
MDA-MB-436
shRNA NT ADAR1
ADAR1
GAPDH
G)
MDA-MB-436
shNT shADAR1
Intensity (Mean Normalized)
H)
MDA-MB-436
shNT shADAR1
Intensity (Mean Normalized)
LPC 16:0
LPC 18:0
PC 32:0
PC 36:1
PC 37:1
PC 38:1
PC 33:2
PC 38:2
PC 40:2
PC 34:3
PC 35:3
PC 44:6
SFA
MUFA
PUFA
I)
PC44:6
Log2 Normalized Intensity
p = 0.0174
shNT shADAR1
J)
MDA-MB-436
shNT shADAR1
Intensity (Mean Normalized)
K)
MDA-MB-436
shNT shADAR1
Intensity (Mean Normalized)
L)
MDA-MB-436
shNT shADAR1
Intensity (Mean Normalized)
M)
shNT shADAR1
Relative Abundance (Fold)
p = 0.929
p = 0.032
p = 0.041
MUFA PUFA XPUFA
Level of unsaturation
N)
PE38:5
Log2 Normalized Intensity
p = 0.0399
shNT shADAR1

**Figure S3. ADAR1 reduction leads to lipid remodeling**

**A)** Concentration of GSH in ADAR1-intact and -deficient MDA-MB-231 cells. n=3. Error bars mark standard deviation. **B)** Ratio of reduced/total GSH to indicate GSH activity in MDA-MB-231 cells. **C)** Light and fluorescent images showed morphology (Bright Field) and ferrous ion detection (FerroOrange) in MDA-MB-231 cells. **D)** Quantification of ferrous ion detection, normalized to cell numbers. n=3. Error bars mark standard error **E)** Significantly altered PC species (red dots) in ADAR1-deficient MDA-MB-231 cells. PUFA PC species are annotated. **F)** Immunoblot analysis showed ADAR1 reduction in MDA-MB-436 cells treated with shADAR1-containing lentivirus. Images are representative of three replicates. GAPDH, loading control. **G)** Relative abundance of all detected PC species in ADAR1-intact (red dots) and -deficient (blue dots) MDA-MB-436 cells. Values were log10 transformed, mean centered, and shown as mean over standard deviation ratio. n=3. **H)** Relative abundance of selected PC species in ADAR1-intact and -deficient MDA-MB-436 cells. **I)** Relative abundance of PC44:6 in MDA-MB-436 cells. **J)** Relative abundance of all detected PC, PE, and PS species in MDA-MB-436 cells, listed in the order of lipid unsaturation. **K)** Relative abundance of PE and PS species in MDA-MB-436 cells, listed in the order of carbon chain. **L)** Relative abundance of PE and PS species in ADAR1-intact and -deficient MDA-MB-436 cells, listed in the order of lipid unsaturation. **M)** Enrichment of PE and PS species with different unsaturation levels (MUFA, 1 double-bond; PUFA, ≥ 2 double-bond; XPUFA, ≥ 5 double-bond) upon ADAR1 knockdown in MDA-MB-436 cells. **N)** Relative abundance of PE38:5 in MDA-MB-436 cells.

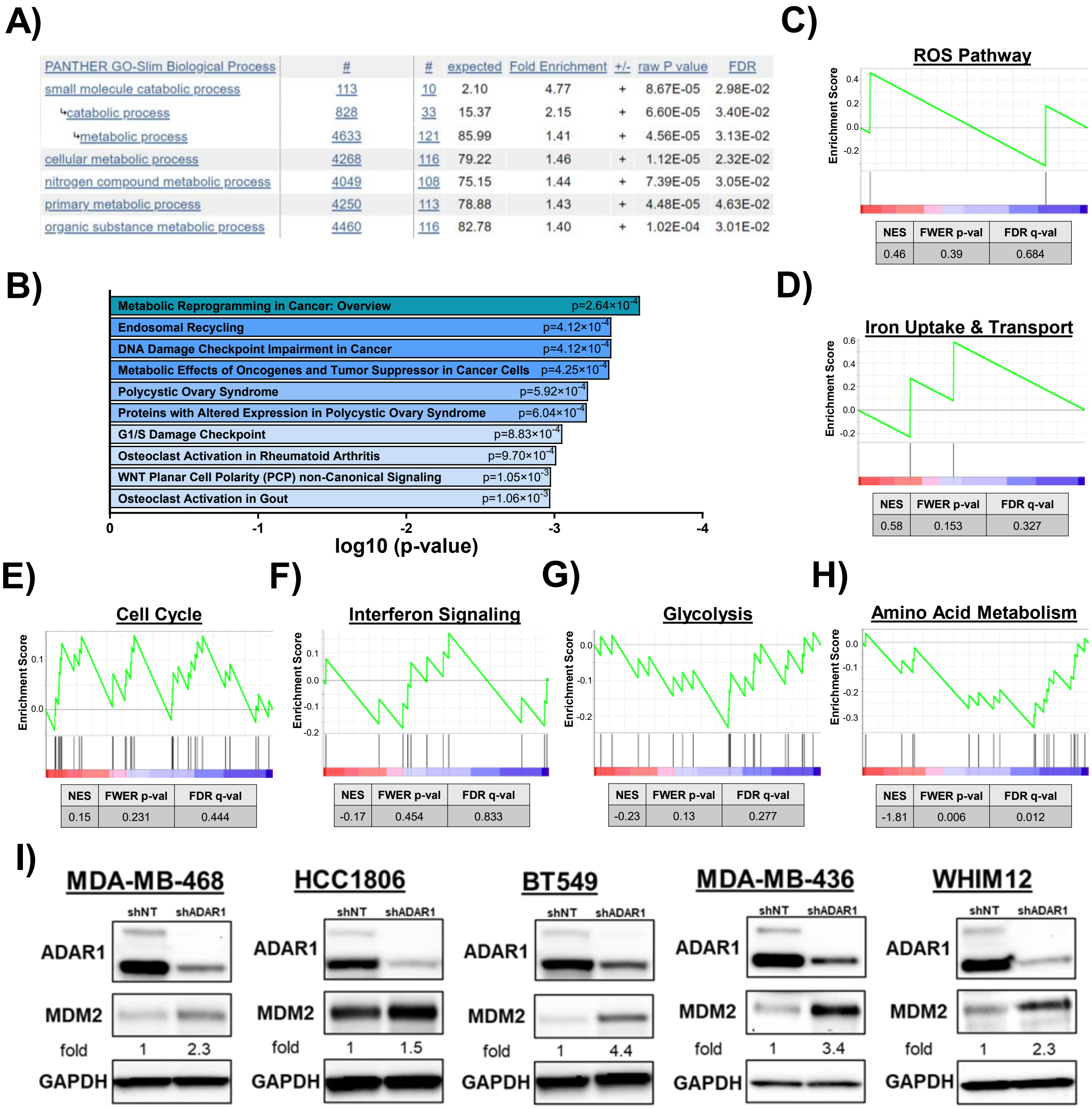
A)
PANTHER GO-Slim Biological Process | # | # | expected | Fold Enrichment | +/- | raw P value | FDR
small molecule catabolic process | 113 | 10 | 2.10 | 4.77 | + | 8.67E-05 | 2.98E-02
catabolic process | 828 | 33 | 15.37 | 2.15 | + | 6.60E-05 | 3.40E-02
metabolic process | 4633 | 121 | 85.99 | 1.41 | + | 4.56E-05 | 3.13E-02
cellular metabolic process | 4268 | 116 | 79.22 | 1.46 | + | 1.12E-05 | 2.32E-02
nitrogen compound metabolic process | 4049 | 108 | 75.15 | 1.44 | + | 7.39E-05 | 3.05E-02
primary metabolic process | 4250 | 113 | 78.88 | 1.43 | + | 4.48E-05 | 4.63E-02
organic substance metabolic process | 4460 | 116 | 82.78 | 1.40 | + | 1.02E-04 | 3.01E-02
B)
Metabolic Reprogramming in Cancer: Overview p=2.64×10-4
Endosomal Recycling p=4.12×10-4
DNA Damage Checkpoint Impairment in Cancer p=4.12×10-4
Metabolic Effects of Oncogenes and Tumor Suppressor in Cancer Cells p=4.25×10-4
Polycystic Ovary Syndrome p=5.92×10-4
Proteins with Altered Expression in Polycystic Ovary Syndrome p=6.04×10-4
G1/S Damage Checkpoint p=8.83×10-4
Osteoclast Activation in Rheumatoid Arthritis p=9.70×10-4
WNT Planar Cell Polarity (PCP) non-Canonical Signaling p=1.05×10-3
Osteoclast Activation in Gout p=1.06×10-3
log10 (p-value)
C)
ROS Pathway
Enrichment Score
NES 0.46 | FWER p-val 0.39 | FDR q-val 0.684
D)
Iron Uptake & Transport
NES 0.58 | FWER p-val 0.153 | FDR q-val 0.327
E)
Cell Cycle
NES 0.15 | FWER p-val 0.231 | FDR q-val 0.444
F)
Interferon Signaling
NES -0.17 | FWER p-val 0.454 | FDR q-val 0.833
G)
Glycolysis
NES -0.23 | FWER p-val 0.13 | FDR q-val 0.277
H)
Amino Acid Metabolism
NES -1.81 | FWER p-val 0.006 | FDR q-val 0.012
I)
MDA-MB-468
HCC1806
BT549
MDA-MB-436
WHIM12
shNT
shADAR1
ADAR1
MDM2
fold
GAPDH
MDA-MB-468 fold 1 2.3
HCC1806 fold 1 1.5
BT549 fold 1 4.4
MDA-MB-436 fold 1 3.4
WHIM12 fold 1 2.3

**Figure S4. Transcriptomic regulation in TNBC cells upon ADAR1 knockdown**

**A)** GeneOntology enrichment analysis for genes significantly altered in ADAR1-deficient TNBC cells, using PANTHER Overrepresentation Fisher's Exact Test. Genes significantly altered in at least 3 cell lines are included. **B)** Significantly altered biological pathways (Elsevier Collection Pathway) in ADAR1-deficient TNBC cells. GSEA enrichment analyses were performed to assess **C)** ROS pathway; **D)** Iron uptake and transport; **E)** Cell cycle; **F)** Interferon signaling; **G)** Glycolysis; and **H)** Amino acid metabolism pathways in ADAR1-deficient MDA-MB-231 cells. **I)** Immunoblot analysis showed MDM2 induction upon ADAR1 knockdown in (left to right) MDA-MB-468, HCC1806, BT549, MDA-MB-436, and WHIM12 cells. Images are representative of three replicates. GAPDH, loading control.

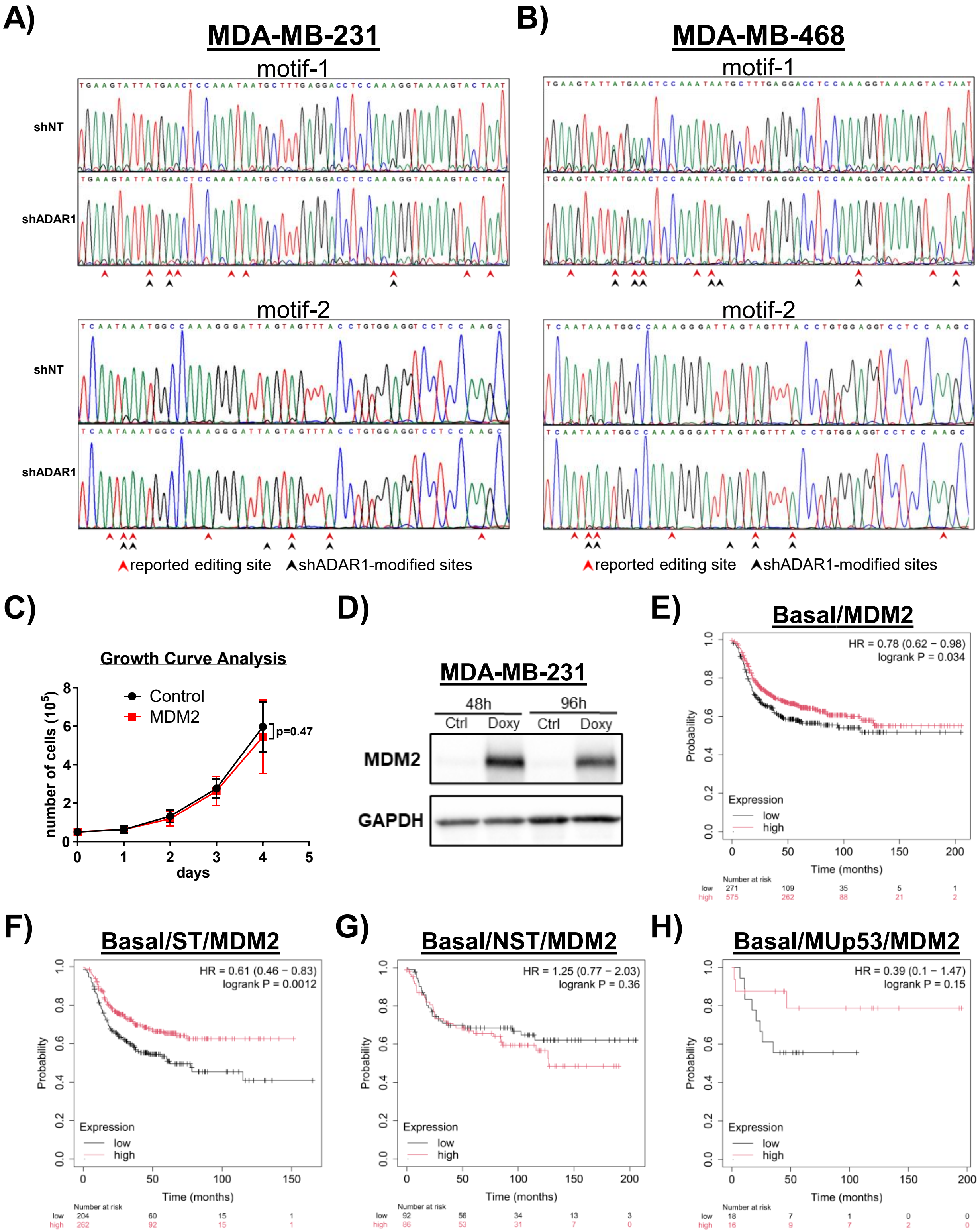
A)
MDA-MB-231
motif-1
shNT
shADAR1
motif-2
shNT
shADAR1
reported editing site
shADAR1-modified sites
B)
MDA-MB-468
motif-1
motif-2
reported editing site
shADAR1-modified sites
C)
Growth Curve Analysis
Control
MDM2
p=0.47
number of cells ($10^5$)
days
D)
MDA-MB-231
48h
96h
Ctrl
Doxy
Ctrl
Doxy
MDM2
GAPDH
E)
Basal/MDM2
HR = 0.78 (0.62 − 0.98)
logrank P = 0.034
Probability
Expression
low
high
Time (months)
Number at risk
low 271 109 35 5 1
high 575 262 88 21 2
F)
Basal/ST/MDM2
HR = 0.61 (0.46 − 0.83)
logrank P = 0.0012
Probability
Expression
low
high
Time (months)
Number at risk
low 204 60 15 1
high 262 92 15 1
G)
Basal/NST/MDM2
HR = 1.25 (0.77 − 2.03)
logrank P = 0.36
Probability
Expression
low
high
Time (months)
Number at risk
low 92 56 34 13 3
high 86 53 31 7 0
H)
Basal/MUp53/MDM2
HR = 0.39 (0.1 − 1.47)
logrank P = 0.15
Probability
Expression
low
high
Time (months)
Number at risk
low 18 7 1 0 0
high 16 9 7 2 0

**Figure S5: ADAR1 regulates MDM2 expression to dictate ferroptosis sensitivity**

Reduced editing of reported ADAR1 editing motifs is shown in ADAR1-deficient **A)** MDA-MB-231 and **B)** MDA-MB-468 cells. A (green) and G (black) reading represent non-edited and edited RNA, respectively. Red arrows mark all reported ADAR1 editing sites in MDM2 3'UTR. Black arrows mark sites with notable reduced editing levels upon ADAR1 knockdown. Sequence chromatograms shown are representative of 3 replicates. **C)** Cell proliferation assay showing that MDM2 overexpression did not alter proliferation of MDA-MB-231 cells. Error bars mark standard deviation, n=3. **D)** Immunoblot analysis showed doxycycline-induced MDM2 overexpression in MDA-MB-231 cells post 48h and 96h doxycycline treatment. Images are representative of three replicates. GAPDH, loading control. MDM2 expression was used to stratify progression-free survival outcomes in **E)** all patients of basal subtype (ESR1-/HER2-) breast cancer, **F)** systematically treated (ST) patients of basal subtype breast cancer, **G)** basal subtype patients who received no systematic treatment (NST), and **H)** patients of basal subtype with confirmed p53 mutations (MUp53). Data from KMplot.com. HR, hazard ratio.

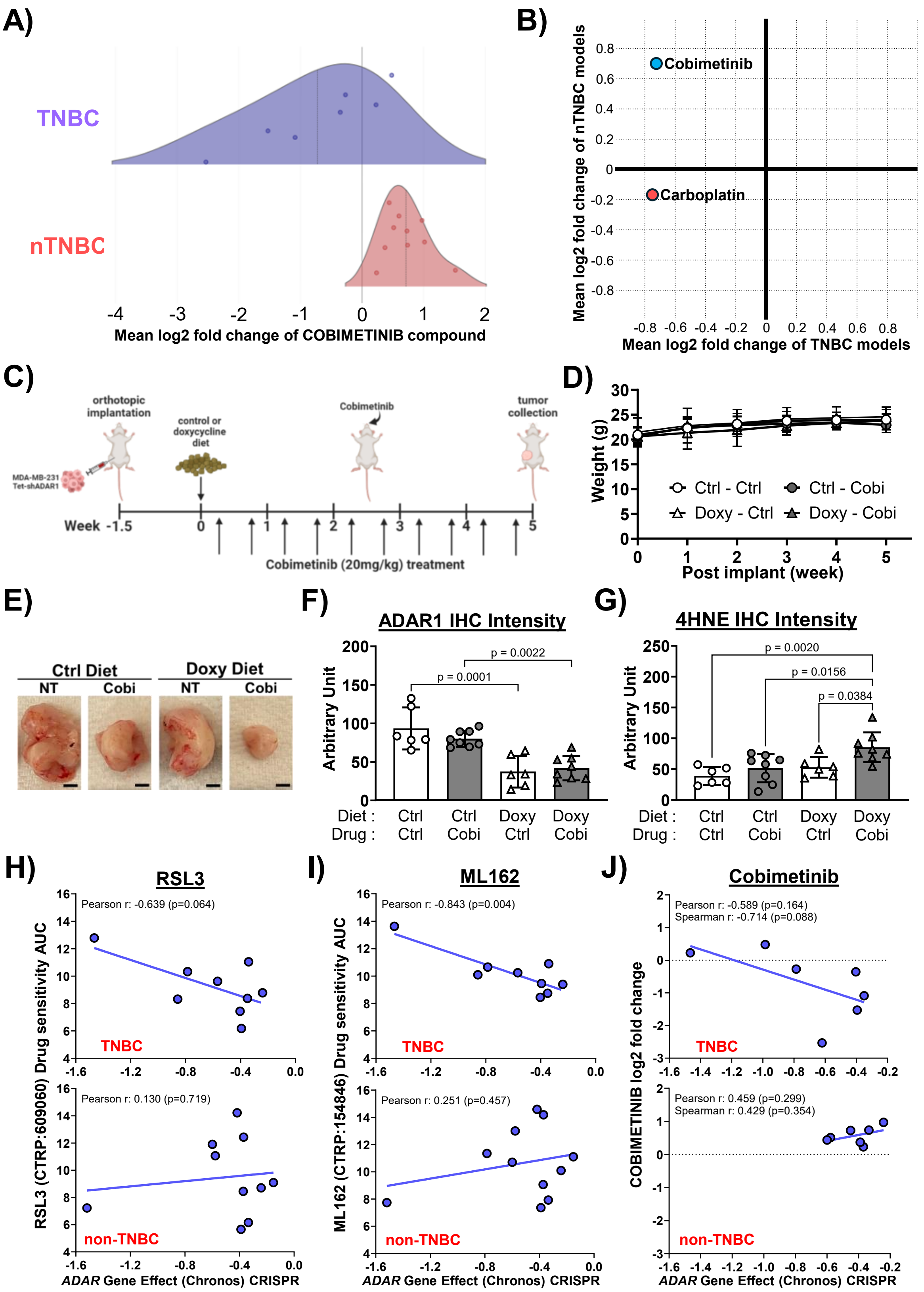
A)
TNBC
nTNBC
-4 -3 -2 -1 0 1 2
Mean log2 fold change of COBIMETINIB compound
B)
Cobimetinib
Carboplatin
Mean log2 fold change of nTNBC models
Mean log2 fold change of TNBC models
C)
orthotopic implantation
control or doxycycline diet
Cobimetinib
tumor collection
MDA-MB-231 Tet-shADAR1
Week -1.5 0 1 2 3 4 5
Cobimetinib (20mg/kg) treatment
D)
Weight (g)
Ctrl - Ctrl
Ctrl - Cobi
Doxy - Ctrl
Doxy - Cobi
Post implant (week)
E)
Ctrl Diet
Doxy Diet
NT
Cobi
NT
Cobi
F)
ADAR1 IHC Intensity
Arbitrary Unit
p = 0.0022
p = 0.0001
Diet : Ctrl Ctrl Doxy Doxy
Drug : Ctrl Cobi Ctrl Cobi
G)
4HNE IHC Intensity
Arbitrary Unit
p = 0.0020
p = 0.0156
p = 0.0384
Diet : Ctrl Ctrl Doxy Doxy
Drug : Ctrl Cobi Ctrl Cobi
H)
RSL3
Pearson r: -0.639 (p=0.064)
TNBC
Pearson r: 0.130 (p=0.719)
non-TNBC
RSL3 (CTRP:609060) Drug sensitivity AUC
ADAR Gene Effect (Chronos) CRISPR
I)
ML162
Pearson r: -0.843 (p=0.004)
TNBC
Pearson r: 0.251 (p=0.457)
non-TNBC
ML162 (CTRP:154846) Drug sensitivity AUC
ADAR Gene Effect (Chronos) CRISPR
J)
Cobimetinib
Pearson r: -0.589 (p=0.164)
Spearman r: -0.714 (p=0.088)
TNBC
Pearson r: 0.459 (p=0.299)
Spearman r: 0.429 (p=0.354)
non-TNBC
COBIMETINIB log2 fold change
ADAR Gene Effect (Chronos) CRISPR

**Figure S6. ADAR1 reduction synergizes with cobimetinib to suppress tumorigenesis**

**A)** Cobimetinib displays superior efficacy against TNBC (n=7) over non-TNBC (n=10) cell lines. Efficacy numbers (PRISM Repurposing Public 24Q2) <0 indicate reduced viability with treatment. Data are extracted from DepMap (depmap.com). **B)** Cobimetinib (TNBC: -0.72; non-TNBC: 0.71) displays superior preference, compared to carboplatin (TNBC: -0.75; non-TNBC: -0.18), against TNBC (n=7) over non-TNBC (n=10) cell lines. **C)** MDA-MB-231 cells (1.5 X $10^5$ per mouse) with Tet-inducible shADAR1 were implanted into mammary fat pads of female mice. Ten days post implantation, half of the mice were switched to doxycycline-containing (625mg/kg) diet, followed by cobimetinib treatments (20mg/kg weight; oral administration twice a week) for 5 weeks. **D)** Progression of average mouse weight. n=5-10 each group. Error bars mark standard deviation**. E)** Representative images of tumors resected after the treatment regimens concluded. Scale bar, 5mm. **F)** Quantification of IHC staining intensity of ADAR1. Error bars mark standard deviation. N=6-8 each group. **G)** Quantification of IHC staining intensity of 4HNE. Error bars mark standard deviation. N=6-8 each group. Drug sensitivity of TNBC, but not non-TNBC, cell lines to **H)** RSL3 (TNBC, n=9; non-TNBC, n=10), **I)** ML162 (TNBC, n=9; non-TNBC, n=11), and **J)** cobimetinib (TNBC, n=7; non-TNBC, n=7) negatively correlate with *ADAR* (encoding ADAR1) dependency (Chronos CRISPR Gene Effect). Lower AUC (RSL3, ML162) and efficacy (cobimetinib) numbers indicate higher drug sensitivity. Lower CRISPR Gene Effect numbers indicate higher *ADAR* dependency. Data are extracted from drug sensitivity (RSL3, ML162: Drug sensitivity AUC, CTD[2]; cobimetinib: PRISM Repurposing Public 24Q2) and gene dependency (Public 24Q2+Score, Chronos) data sets from DepMap (depmap.org).
